# Promoting intestinal healing by exclusive enteral nutrition with TGF-β in a mouse model of colitis

**DOI:** 10.1101/2023.08.31.555691

**Authors:** Kawthar Boumessid, Vickie Lacroix, Ekaterina Ovtchinnikova, Muriel Quaranta Nicaise, Maryline Roy, Anne Dumay, Sophie Thenet, Marie Carriere, Emmanuel Mas, Frédérick Barreau

**Author notes:** co-corresponding : Addresses for correspondence: Pr. Emmanuel Mas, MD, PhD, Gastroenterology, Hepatology, Nutrition, Diabetology and Hereditary Metabolic Diseases, Unit 330, avenue de Grande-Bretagne, TSA 70034, 31059 Toulouse cedex 9, France; Dr. Frédérick Barreau, Institut de Recherche en Santé Digestive (IRSD), INSERM U1220, Purpan Hospital, CS60039, 31024 Toulouse, cedex 03, France. co-last authors (Emmanuel Mas, Frédérick Barreau). **Author contributions:** study concept and design : FB and EM; acquisition of data : KB, VL, EK, MQN, MR, AD, FB; analysis and interpretation of data: KB, VL, AD, ST, MC, EM and FB; drafting of the manuscript: KB, AD, ST, MC, EM and FB; critical revision of the manuscript for important intellectual content: KB, AD, ST, MC, EM and FB; statistical analysis: KB, AD, FB and EM; obtained funding : EM and FB;, technical, or material support:MC and ST; study supervision: EM and FB.

## Abstract

**Background and aims:** Exclusive Enteral Nutrition (EEN) is the first line of treatment for pediatric Crohn’s disease (CD), but its mechanisms of action remain poorly understood. We studied EEN nutritional composition and TGF-β effect in a mouse model of colitis, as well as the role of intestinal microbiota.

**Methods:** Mice were treated with Dextran Sulfate Sodium (DSS) during 5 days to induce colitis, until the inflammatory peak (day 7) or gut restitution (day 14). After DSS treatment, some of them received EEN formula such as Modulen IBD® (DM mice) or Infatrini Peptisorb® (DINF mice), with TGF-β supplementation or neutralization, and clinical inflammation was evaluated. After sacrifice, macroscopic and microscopic inflammation were analyzed, as well as intestinal permeability (IP). The composition of mucosal colonic microbiota was analyzed and fecal microbiota transplantation was performed to evaluate its capacity to mediate anti-inflammatory and pro-regenerative effect. Colonic crypts from DSS and EEN mice were cultured as 3D organoids and cellular properties were analyzed.

**Results:** DSS mice developed colitis, as evidenced by the weight loss and clinical inflammation. It was accompanied by macroscopic inflammation such as colon thickness and edemas, and an elevated IP. In contrast, EEN mice with TGF-β formula present faster weight recovery and decreased inflammatory parameters, with a normalized IP, suggesting gut restitution and functionality. These functional improvements were not obtained for EEN mice without TGF-β formula. Moreover, EEN with Modulen IBD® (DM mice) modified the microbiota in comparison to DSS condition and attenuated inflammation. In addition, the organoids from DM mice colonic crypts treated had an enhanced survival, and re-epithelialization capacity.

**Conclusions:** Both EEN formula have anti-inflammatory properties, certainly by the nutritional composition. However, TGF-β plays a significant role in intestinal restitution and restoring barrier function. These beneficial effects are partly mediated by the microbiota to maintain gut homeostasis.

## INTRODUCTION

Crohn’s disease (CD), a type of Inflammatory Bowel Disease (IBD) along with Ulcerative colitis (UC), represents a worldwide health issue since its incidence is continuously increasing. IBD are characterized by chronic relapsing inflammation of the digestive tract. Indeed, CD can affect the entire digestive tract, from the mouth to the anus with discontinuous and deep lesions. It is due to both genetic and environmental factors. Among the genetic factors, mutations in *NOD2* (Nucleotide-binding Oligomerization Domain-containing protein 2) gene were the first to be characterized, and have been associated with the highest risk of developing CD.^1^ This gene encodes for the NOD2 protein, an intracellular receptor that recognizes muramyldipeptide, a fragment of the peptidoglycan presents in both Gram-positive and Gram-negative bacteria, described to regulate the homeostasis of the digestive mucosa.^1, 2^ Concerning the environmental factors, only smoking, antibiotics, and meat consumption are associated with CD development.

The pathophysiology of CD is described to be the result of an abnormal immune response associated with an altered intestinal microbiota, called dysbiosis.^2, 3, 4^ Although the causal link(s) between these alterations is not known, a barrier defect of the intestinal mucosa is involved in the genesis and maintenance of this abnormal communication, leading to inflammation and epithelium alterations.^5^ In addition, intestinal permeability (IP) is increased, along with an alteration of the tight junctions’ proteins. Thus, in IBD, the four compartments of the barrier function of the intestinal mucosa, i.e. the microbiota, the mucus layer with antimicrobial peptides, the intestinal epithelium, and the immune system are altered. The integrity of the intestinal epithelium is crucial and is ensured by intestinal stem cells (ISC), which allow the renewal of the intestinal epithelium in 3 to 5 days in mice.^6^ The ISCs are endowed with self-renewal and multipotency capacities, by generating progenitors that will differentiate into mature and functional cells. In IBD, a default of the epithelial renewal involving an excessive immune stimulation has been documented.

The main treatments for CD are anti-inflammatory drugs, immune regulators, and biologics, targeting the immune system or the inflammatory process.^7^ Although these treatments improve symptoms, they are not sufficient to avoid a relapse. Another possible treatment that has proved its clinical efficiency is the exclusive enteral nutrition (EEN), with clinical studies reporting up to 86% of clinical remission.^7, 8^ According to the newest recommendations,^9^ EEN is the first induction treatment for pediatric CD in case of non-stricturing or penetrating inflammation, and without severe growth delay. Besides its clinical efficiency, EEN permits mucosal healing. By definition, mucosal healing is the absence of ulcerations and lesions observed by endoscopy, suggesting a mucosal regeneration.^7^ By far, mucosal healing represents the best criteria for long-term clinical remission, which makes it a therapeutic objective for CD. While many mechanisms could explain these results,^7^ the EEN formula compositions and nutritional matrix could play a crucial role. Nutritional literature has already discussed these two characteristics, as well as their consequences on physiology and pathophysiology.^10^ In the case of IBD, decreasing intestinal inflammation, and permeability, as well as restoring the gut barrier would be hypothetical.

Many mediators regulate the epithelium renewal, among which is the Transforming growth factor β (TGF-β).^11^ TGF-β is a cytokine produced by immune cells, intestinal epithelial cells, and fibroblasts, and is involved in the first phase of mucosal healing. The first phase of this healing also called “restitution” is based on the redistribution of existing cells to fill in the ulcerated areas without epithelial cells. In addition to its role in epithelial wound healing, TGF-β is also described to maintain the integrity of tight junctions, prevent Goblet cells’ depletion, and is involved in immune regulation. The TGF-β production and/or signaling are dysregulated in IBD, which alters the effects of TGF-β. Thus, in this study, we have investigated the mechanisms by which a polymeric formula containing TGF-β alleviated the colitis and strengthened the epithelial renewal.

## MATERIAL AND METHODS

### Mice

C57BL/6Nrj wild-type and *Nod2^KO^* male mice of twelve weeks of age, from CREFRE or Janvier Labs, were hosted in the animal facility of Purpan (Toulouse, France).^12^ Mice were treated with dextran sulfate sodium (DSS) at 3% during 5 days to induce a colitis, and then DSS was replaced with regular water. Progression of the disease was monitored daily until the 14^th^ day. Instead of regular food, other mice received Modulen IBD® *(Nestlé)* or Infatrini Peptisorb® *(Nutricia) ad libitum* from day 5 to day 7 or 14. For TGF-β neutralization, 1D11 antibody *(ThermoFischer Scientific)* was added to Modulen IBD® solution during the treatment. For TGF-β signaling inhibition, SB431542 *(Selleckchem)* was diluted in DMSO, PEG300 and distilled H2O following the manufacturers’ instructions, and administrated *via i.p* every day from day 4 to day 7, then every two days from day 7 to day 14. Finally, Recombinant TGF-β2 proprotein expressed in mammalian cell *(MBS954812, MyBiosource)* was added to Infatrini Peptisorb® at 10 ng/mL and administrated to mice *ad libitum*. TGF-β2 concentrations were measured by ELISA kit CSB-E14209B *(Cusabio)* according to their protocol. Animal experimentations were approved by the ethical committee, and performed under the protocols “APAFIS#28278-2020111810056802, and 2020120418538745”.

### Clinical observations

To assess colitis severity, mice body weight was monitored daily in order to establish a disease activity index (DAI) adapted from Van der Slui et al.^13^ We attributed points for weight loss as following: 1 point if weight loss was between >5% and <10%, 2 points if ⍰10% and <15%, 3 points if ⍰15% and <20%, and 4 points if ⍰20%. Presence of blood in stools or around anus corresponded to 1 point. Non-formed stools corresponded to 1 point, mild diarrhea to 2 points, and severe diarrhea to 3 points. Physical appearance was also observed: 1 point for bristling fur, 2 points if hunched back, 3 points if both, and 4 points if the mice was lethargic.

### Macroscopic analysis

A macroscopic inflammation score was determined by analyzing colon appearance, taking into account edema length (1 point if < 1 cm; 1.5 if ⍰1 cm and < 2cm; 2 points if ⍰2cm), and colon wall thickness measuring with a caliper (0 points if <0.3mm, 1 point if ⍰0.3 and < 0.45mm, 1.5 points if ⍰0.45 and <0.55mm, and 2 points if ⍰0.55mm). Colon length (cm) was also monitored.

### Histological analysis

To establish histological analysis, colon rolls were established. The colon was enrolled, fixed in formol overnight, then in ethanol 70% at 4°C. After inclusion in paraffin, sections of 5 µm were made for hematoxylin / eosin coloration. Photos were acquired with Panoramic Scan Flash III, and analyzed with CaseViewer. A minimum of 49 crypts by sample, from the entire sample, was measured, and their cell number was counted. For each condition, two different samples were analyzed. The histological score was adapted from the literature,^14^ and points were attributed as: inflammatory cell infiltrate extend (Mucosal: 1, Mucosal and submucosal: 2, Mucosal, submucosal and transmural: 3), epithelial changes (Hyperplasia (minimal <25%: 1, Mild 25-35%: 2, Moderate 36-50%: 3, Marked > 51%: 4), Goblet cell loss (minimal <20%: 1, Mild 21-35%: 2, Moderate 36-50%: 3, Marked > 50%: 4), Cryptitis: 2 to 3, Crypt abscesses: 3 to 5, Erosion: 1 to 4), mucosal architecture (Ulceration : 3 to 5, Irregular crypts: 4 to 5, Crypt loss: 4 to 5).

### Paracellular permeability measurement

Distal colonic samples were mounted in duplicates in Ussing chambers with 37° Ringer solution.^15^ The mucosal to serosal flux was measured with 4 kDa FITC-Dextran to assess the paracellular permeability.^16^

### Colonic organoid culture

After sacrifice, colons were opened longitudinally and washed in PBS without calcium and magnesium, then transferred to a dissociation medium (PBS, EDTA 9 mM, DTT 3 mM, Y27632 10 µM). Organoid culture is described in supplementary material.

### RNA extraction, reverse transcription and quantitative PCR

RNA from colonic crypts was extracted by Trizol reagent.^17^ RNA extraction, reverse transcription and PCR methodologies are described in supplementary material.

### DNA methylation

The levels of 5-methyl deoxycytosine (5-Me-dC) were measured by high-performance liquid chromatography–mass spectrometry (HPLC-MS/MS). DNA methylation is describd in supplementary material.

### Microbiota sequencing

DNA was extracted from scraped colonic samples using a QIAamp PowerFecalPro DNA kit (*Qiagen, Hilden, MD*), following the manufacturer’s instructions.^18^ DNA extracts were quantified on a Qubit4 fluorimeter using the dsDNA HS Assay Kit (*Life Technologies; USA*) and were stored at −20°C before further analysis. Microbiota sequencing is described in supplementary material.

### Fecal microbiota transplantation

Bedding of control mice or mice treated with DSS and DSS + Modulen IBD® were collected between days 10 and 14, in order to transfer it to C57BL/6Nrj wild-type germ-free mice. After 4 weeks of adaptation and to ensure that the transfer of microbiota was stable,^3^ 3% DSS cycle was performed, and same analyses described above were done.

### Statistical analysis

Statistical analysis was done with GraphPad Prism version 9 (*Graphpad software, San Diego, CA, USA*). For each data, distribution was analyzed. For bodyweight, and DAI monitoring t-test or Mann-Whitney was done. Significance of beta-diversity metrics was assessed by PERMANOVA test (Permutational Multivariate Analysis of Variance Using Distance Matrices). For the bacterial abundance, alpha-diversity and other data ANOVA or Kruskall-Wallis, with multiple comparison tests, was done. All data are represented in mean ± SEM. *P-values* are: * p<0.05; ** p<0.01; ***p<0.001; ****p<0.0001.

## RESULTS

### EEN with Modulen IBD® has anti-inflammatory properties, and promotes intestinal healing in a mouse model of CD

The first step consisted of developing an experimental inflammatory model reproducing the anti-inflammatory properties of EEN, as reported in clinical studies and by our data (Supplementary Figure 1). For this purpose, clinical and macroscopic inflammatory parameters were monitored in DSS mice treated or not with Modulen IBD® (DM). DSS mice lose weight from day 7 to day 11, and exhibit an elevated DAI, evidencing colitis development (Figures 1 A and B). Moreover, macroscopic inflammation is determined with colonic shortening (Figure 1C) and thickening (Figure 1D), accompanied by a high macroscopic inflammatory score, at both sacrifice times, day 7 and day 14 (Figures 1 C-E). DM decreases colitis-induced weight loss significantly, DAI, macroscopic inflammation score, and edema score (Figures 1A-F). DM decreases colon thickness significantly at day 14, whereas there is no significant statistical difference at day 7 (Figure 1D). At the microscopic level, DSS induces a crypt shortening, as well as a decrease of cell number per crypt (Figures 1G and H). It also increases pro-inflammatory cytokine CxCl1/KC relative expression at both day 7 and day 14 of sacrifice, which is improved by DM at day 7 (Figure 1I). The histological score is attenuated by DM, specifically crypt irregularities, hyperplasia and erosions (Supplementary Figure 2). DM raises the total number of cells per crypt, and the crypt length at day 7 but not at day 14 (Figures 1G and H). The macroscopic and microscopic results evidence an improvement of epithelial restitution and barrier function. To assess this last one, and mucosal functionality, paracellular permeability was studied by Ussing chambers. DM normalizes the high colonic permeability induced by DSS (Figure 1J). These results support the anti-inflammatory properties of DM in our mouse model of colitis. Furthermore, similar results were obtained with a mouse model of genetic predisposition to CD (*Nod2^−/−^*) (Supplementary Figure 3). Indeed, EEN with Modulen IBD® decreases weight loss, and DAI, as well as macroscopic inflammation. In addition, restoration of the intestinal barrier may underlie a mucosal regeneration process in WT and *Nod2*^−/−^ mice.

**Figure 1.**
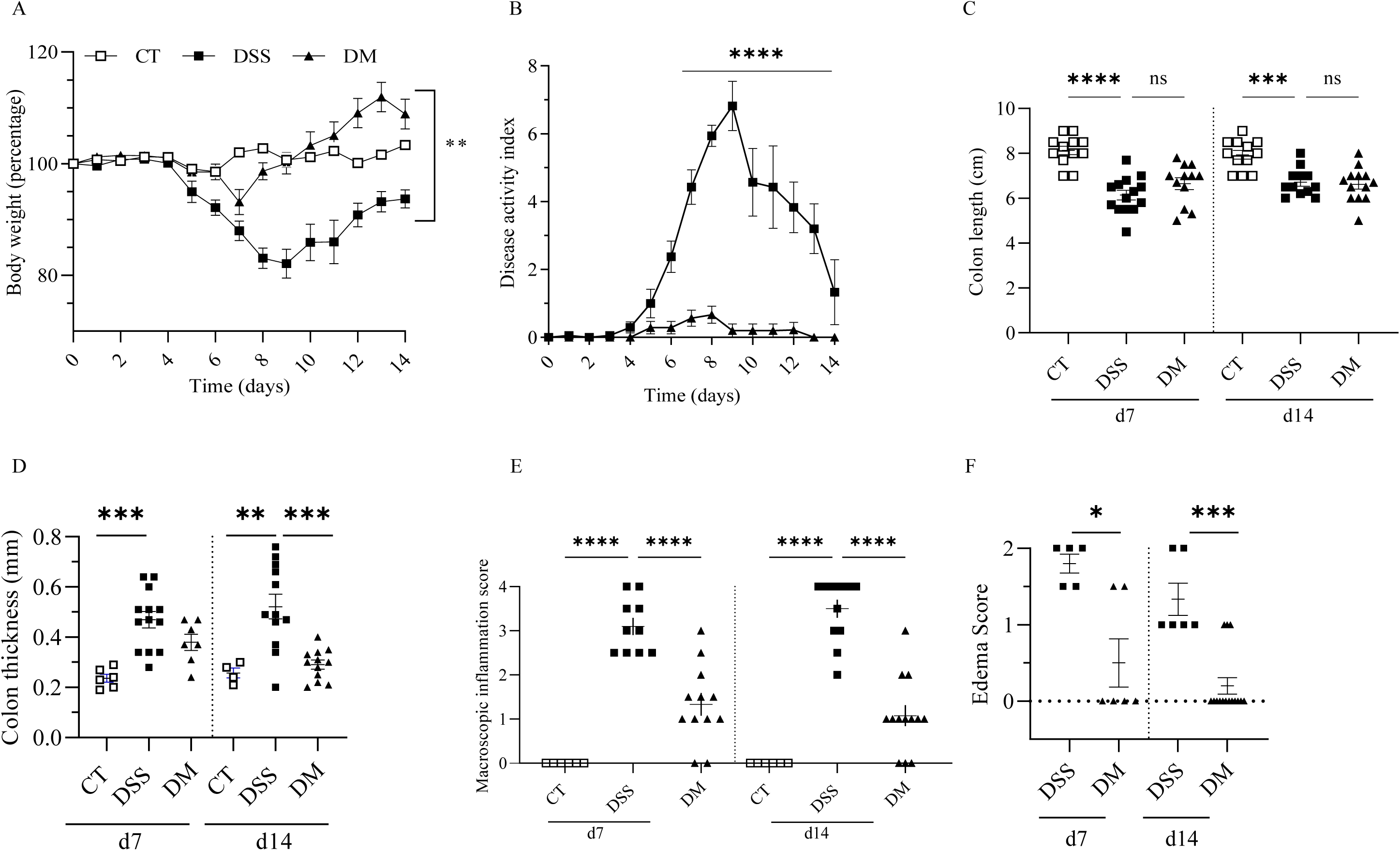
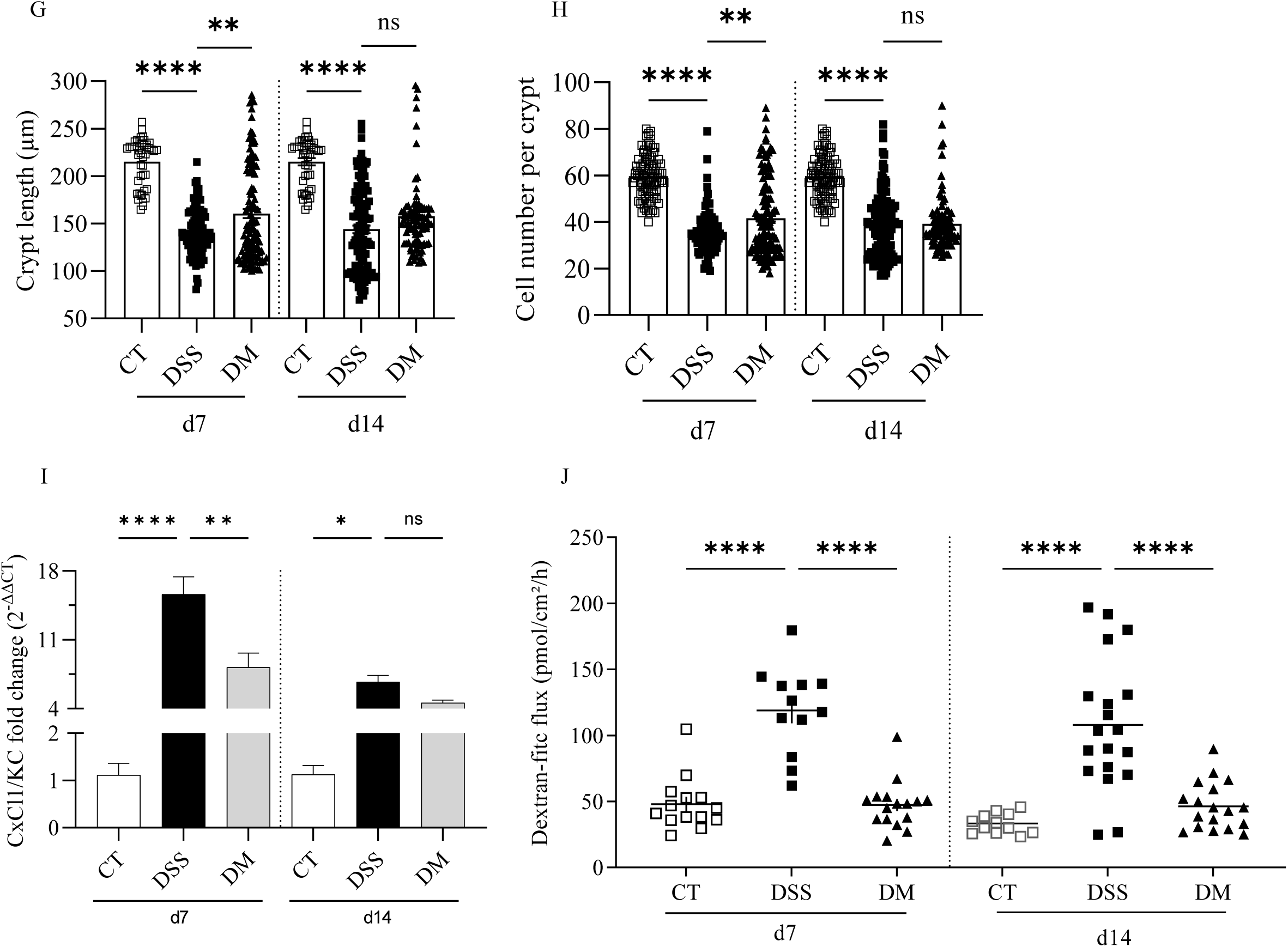
EEN with Modulen IBD® has anti-inflammatory properties, and promotes intestinal healing in a mouse model of colitis. **A:** bodyweight monitoring of CT, DSS and DM treated mice (legend at the top). **B:** DAI (disease activity index) during DSS procedure, and DM mice (same legend as A). **C:** Colon length at day 7 and day 14 expressed in cm. **D:** colon thickness at day 7 and day 14 expressed in mm. **E**: macroscopic inflammatory score at day 7 and day 14. **F:** edema scoring of DSS and DM mice at days 7 and 14. **G:** Crypt length measured in µm at day 7 (minimum of 44 crypts per sample, two samples per condition). **H:** total cell number per crypt at day 7 (minimum of 44 crypts per sample, two samples per condition). **I:** mRNA relative expression of CxCl1/KC from crypts of CT, DSS, and DM mice (n>4). **J:** FITC-Dextran flux representing intestinal permeability performed in duplicate, each point equivalent to a duplicate. Results are expressed as mean ± SEM; at least n>7 mice per group. *P-values*: * p<0.05; ** p<0.01; ***p<0.001; ****p<0.0001 vs. DSS group.

### Nutritional matrix and TGF-β2 of EEN formula are necessary to maintain intestinal homeostasis in a mouse model of CD

The EEN mechanisms are still poorly understood and several EEN components could be involved in their beneficial effects on intestinal health. The nutritional matrix, as well as TGF-β2 contained in some formula (Modulen IBD® and Santactiv Digest®) could play an important role that we studied (Supplementary Table 2). The concentrations of TGF-β2 in formula were quantified by ELISA and showed that Modulen IBD® contained 8 ng/ml of TGF-β2 whereas it was undetected in Infratrini Peptisorb® (Supplementary Figure 4). Both EEN formula used in this study (Infatrini Peptisorb® (DINF) and Modulen IBD® (DM)) decreased clinical parameters of inflammation (Figure 2). DM tends to have a greater weight recuperation than DINF, but the difference is not significant (Figure 2A). The supplementation of Infatrini Peptisorb® (DIT) and neutralization in Modulen IBD® (DMN) with TGF-β2 do not affect weight loss, DAI or colonic length (Figures 2A-C). In contrast, only the DM formula with higher concentration of TGF-β2 decreases the colon thickness at day 14 (Figure 2D), macroscopic scoring (Figure 2E), edema score (Figure 2F) and restores a normal intestinal permeability (Figure 2G). Some of these beneficial impacts are lost when TGF-β2 is neutralized (DMN), including macroscopic inflammatory score, edema score, and intestinal permeability (Figures 2 E-F). On the other hand, supplementation of Infatrini Peptisorb® with TGF-β2 (DIT) tends to decrease macroscopic inflammatory score and edema scoring, and even completely abolished in some mice at day 14, as observed for DM (Figures 2E and F). Regarding colonic permeability, formula without or with less TGF-β2 (DMN/DINF) do not mitigate hyperpermeability induced by DSS, whereas its supplementation (DIT) normalized it (Figure 2G). These results suggest that both the nutritional matrix and TGF-β play a role in the anti-inflammatory properties of EEN. However, TGF-β2 seems to play a major role in the functionality of the epithelium by restoring gut permeability. Inhibition of TGF-β signaling pathway during EEN (DMI) confirms these results (Supplementary Figure 5). This inhibition abrogated anti-inflammatories properties of EEN, more specifically weight recovery, DAI, edema and macroscopic scores. Also, IP remained increased similarly to DSS condition.

**Figure 2.**
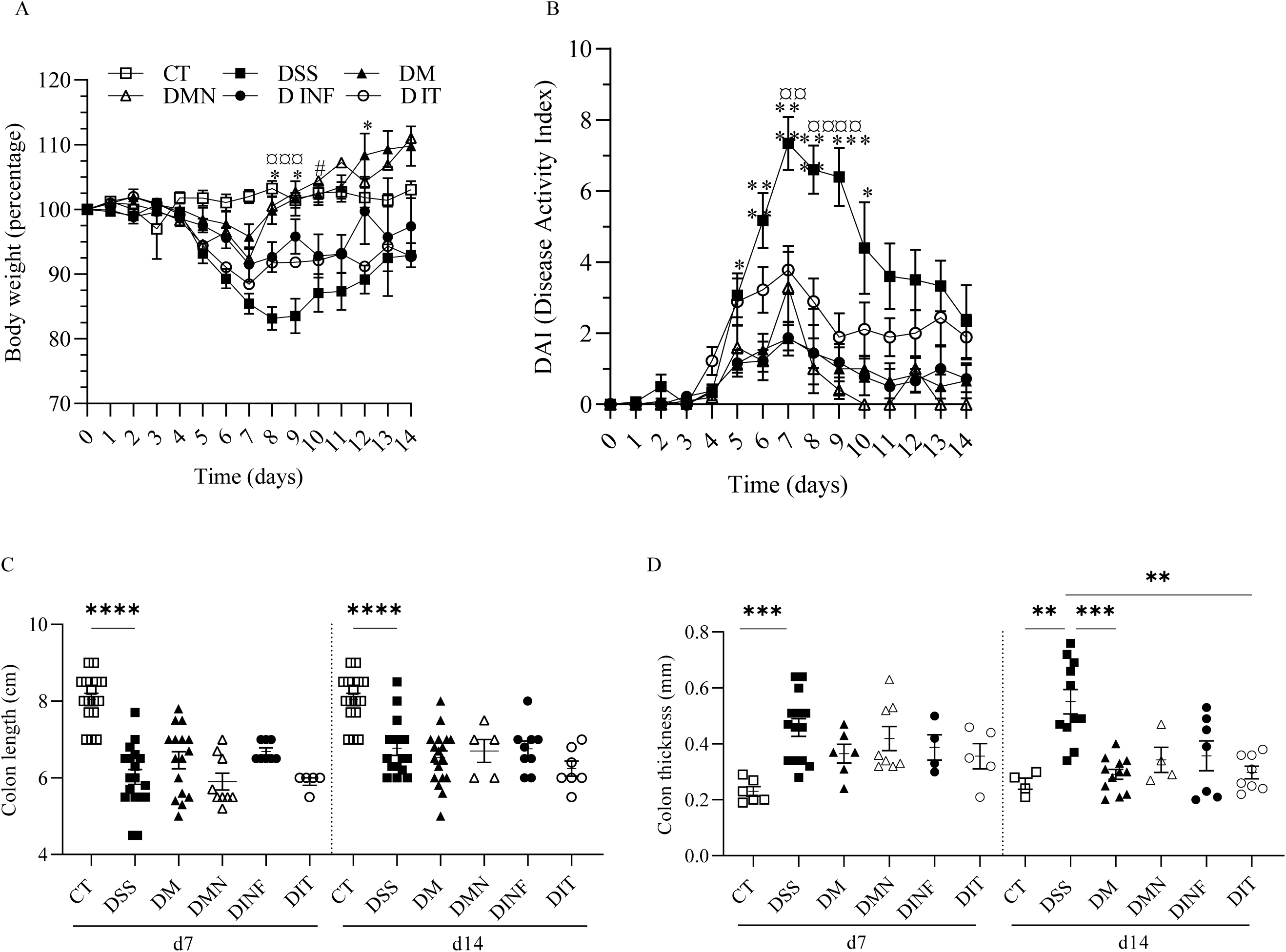
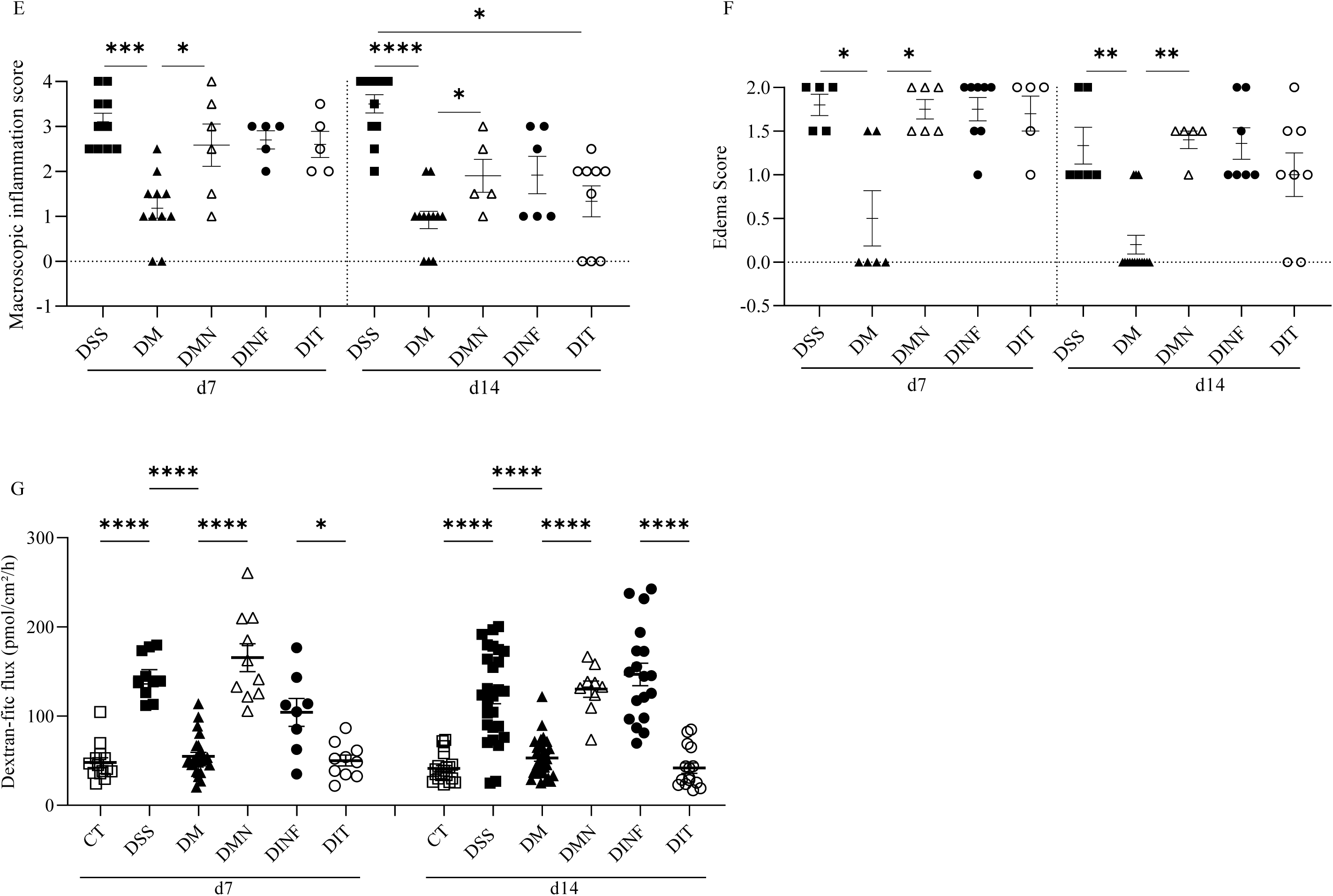
Nutritional matrix and TGF-β of EEN formula are necessary to restore intestinal homeostasis in a mouse model of colitis. **A:** bodyweight evolution of CT (control), DSS, DM (DSS + Modulen IBD®), DMN (DSS + Modulen IBD® TGF-β neutralization), DINF (DSS + Infatrini Peptisorb®), DIT (DSS + Infatrini Peptisorb® + TGF-β) (legend at the top) (n >10). Statistical analysis: * (DINF vs DSS), # (DINF vs DM), ¤ (DIT vs DM). **B:** DAI (Disease Activity Index) during experimentation (same legend as A). **C:** colon length in cm at d7 and d14. **D:** colon thickness in mm at d7 and d14. **E**: macroscopic inflammation score at d7 and d14. **F:** edema scoring at d7 and d14. **G:** intestinal permeability performed in duplicate, each point equivalent to a duplicate. Data are expressed as mean ± SEM; at least >5 mice per group. *P-values*: * p<0.05; ** p<0.01; ***p<0.001; ****p<0.0001.

### EEN shapes the intestinal microbiota

Since microbiota is known to be modified by nutrients and is involved in gut homeostasis, bacteria adherent to the colonic mucosa were investigated by 16S rDNA sequencing (Figure 3). DSS-treated mice microbiota presents low alpha diversity, as shown by the Shannon index, at both day 7 and day 14 (Figure 3A). During inflammation at day 7, beta-diversity analysis shows different composition between DM and DSS groups (Figure 3B). At phyla levels, in comparison with DSS, DM treatment induces a decrease of *Firmicutes* and an increase of *Bacteroidota, Proteobacteria* and *Desulfobacterota* (Figures 3C and Supplementary Figure 6). At day 14, according to beta-diversity shown in Figure 3B, DSS or DM treated mice exhibited a significant different microbial composition in comparison to control mice. An increase of the phylum *Cyanobacteria* is observed in DSS-treated mice in comparison to control, and an increase of *Proteobacteria* by Modulen IBD® treatment (DM condition compared to DSS) (Figure 3C and Supplementary Figure 6). A differential analysis (DESeq2) was applied to study the details of changes in mucosal microbiome composition (Figure 3D). At day 7, 10 marker taxa belonging to *Firmicutes* and 1 to *Desulfobacterota* phyla were identified, in comparison of DM to DSS groups (Figure 3D). In response to Modulen IBD® treatment, 6 main genus are over-represented with a log2 fold change between 2.4 and 21.5, while 5 genus are reduced with a log2 fold change between 4.2 and 7.2 (Figure 3D). The mostly increased genus by Modulen IBD® treatment is *Lachnospiraceae_NK4A136_*(cluster_244: 21.5 log2 fold change in DM vs DSS), which is drastically decreased by DSS treatment (cluster_244: 16.8 log2 fold change in DSS vs CT). DM treatment also increased another species belonging to the *Lachnospiraceae_NK4A136_*group (cluster_90: 5.17 log2 fold change in DM vs DSS, Figure 3D). At the opposite, the both increased genus clusters 10 (*Lactobacillaceae / Lactobacillus*) and 18 (*Enterococcaceae / Enterococcus*) by DSS were reduced by Modulen IBD® treatment by 4.2 and 7.2 log2 fold change respectively (Figure 3 D). Finally, Modulen IBD® treatment also reduced species belonging to the *Lachnospiraceae / Roseburia* genus (clusters 103, 112 and 150: 6.3, 6.2, 5.2 log2 fold change in DM vs DSS, Figure 3 D).

**Figure 3.**
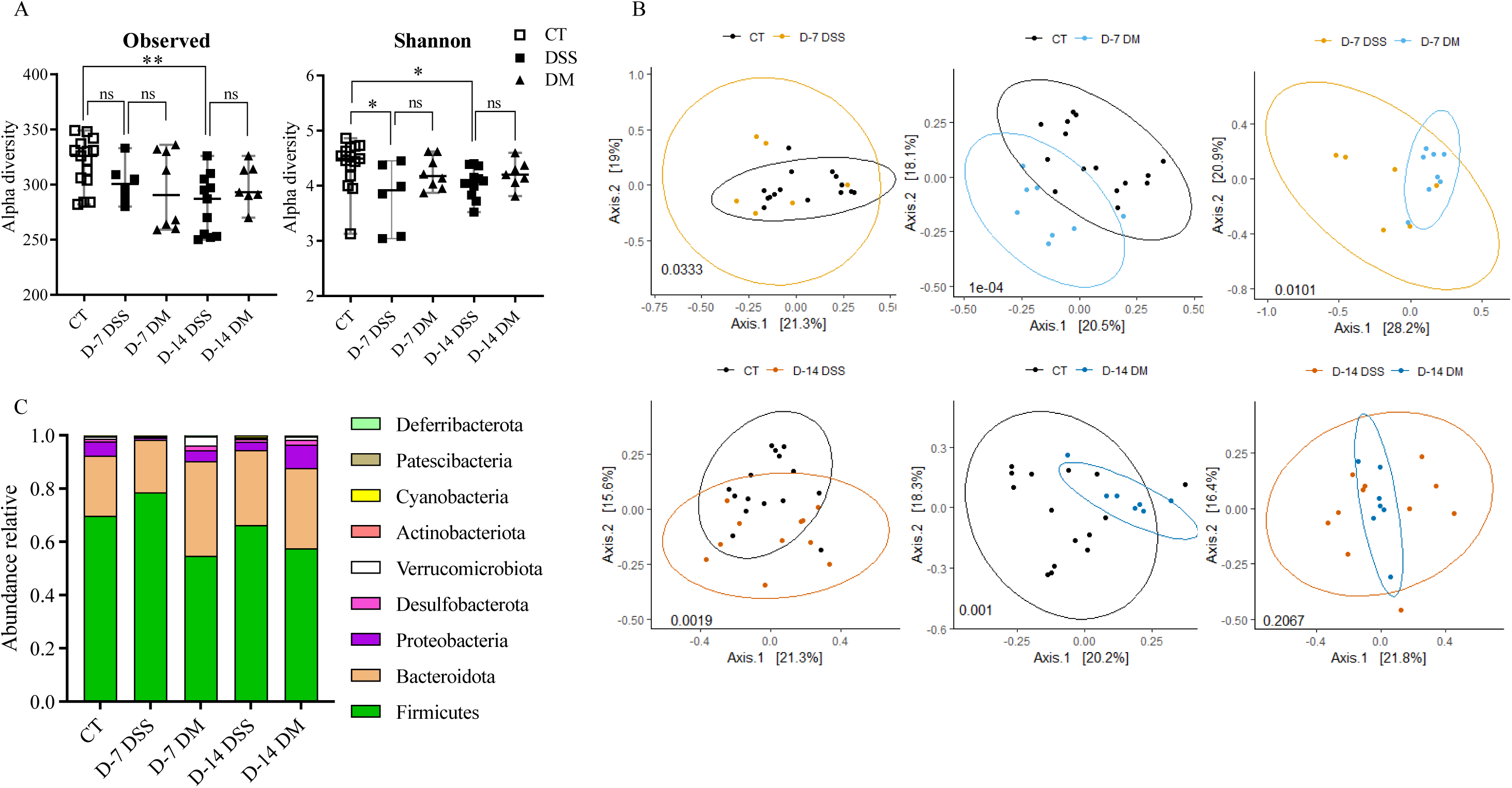
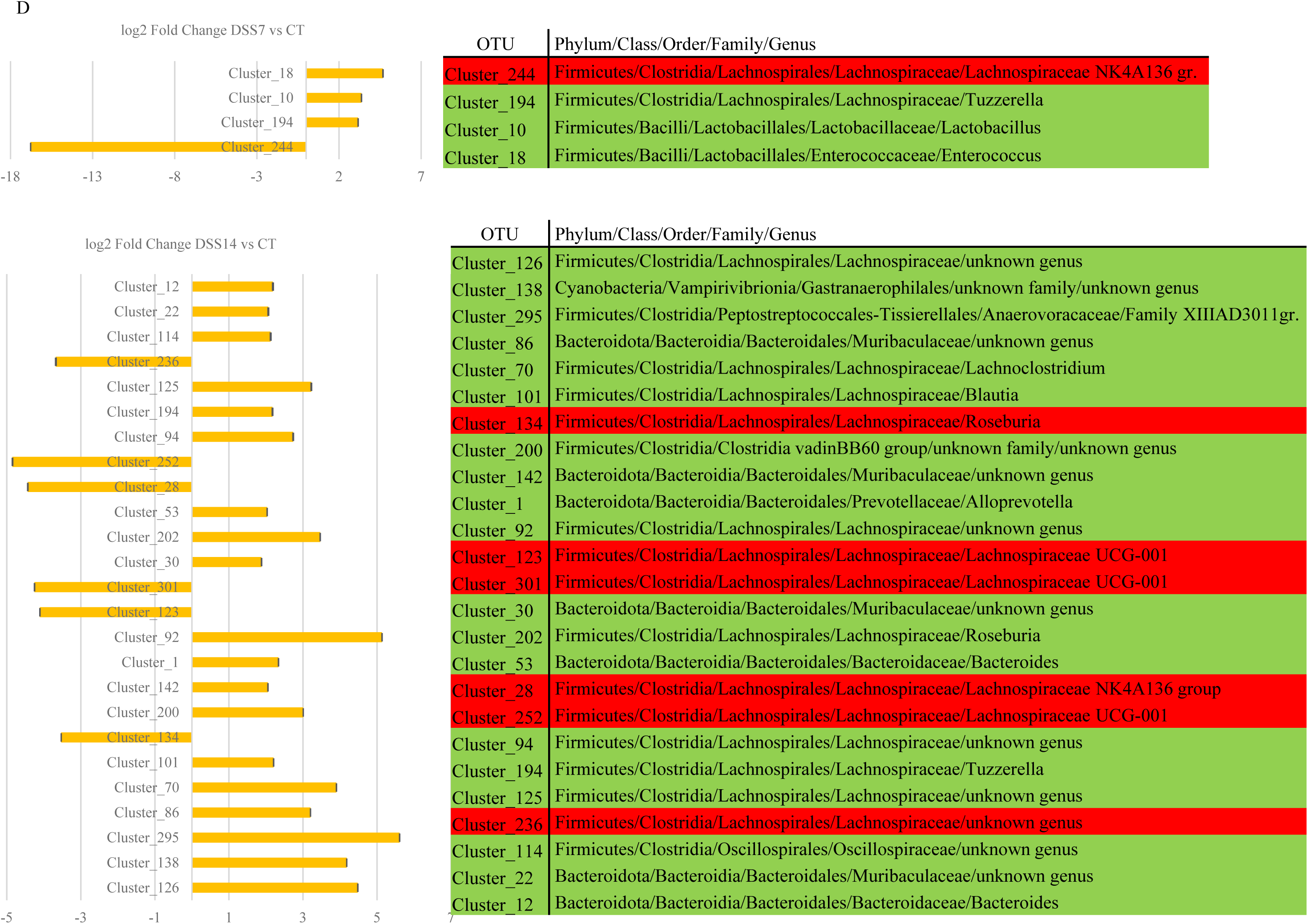
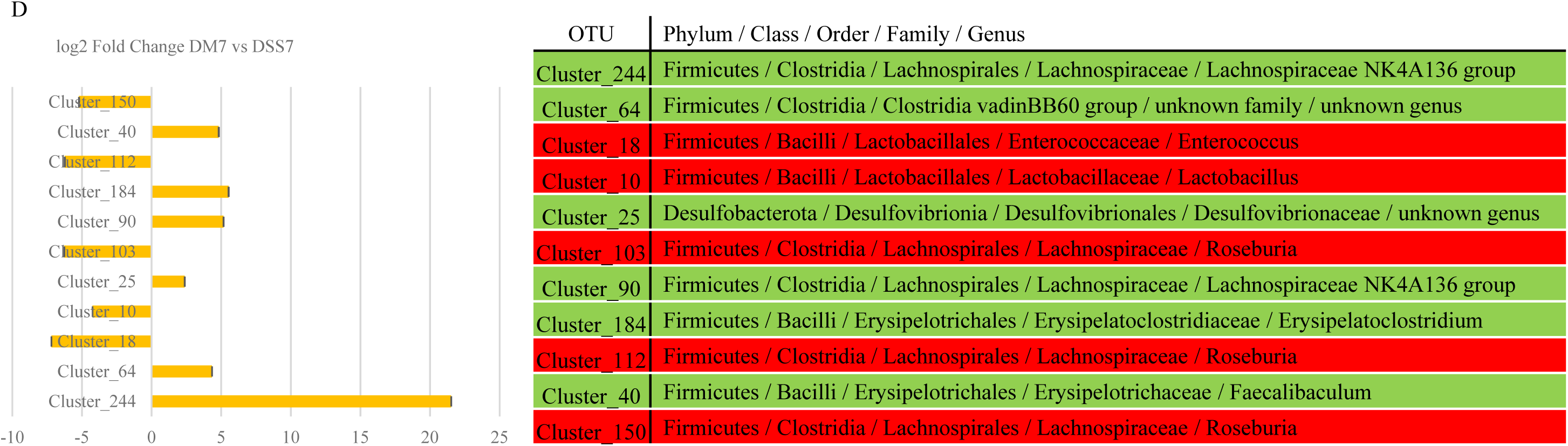
EEN shapes the intestinal microbiota. **A:** alpha-diversity analyzed by observed OTUs richness and Shannon index. Data are expressed as median with minimum/maximum ranges. **B:** Principal Coordinates Analyses (PcoA) of bacterial beta-diversity in CT, DSS or DM mice at day 7 or day14, using Bray-Curtis distance plots. *P-value* were determined by PERMANOVA test. **C:** Relative abundance of phyla in each group. **D:** DESeq2 differential abundance analysis. Differentially abundant taxa between groups are represented by log2 fold change with a *P-value* < 0.01.

Such as the EEN with Modulen IBD® (DM group), Infatrini Peptisorb® formula (DINF group) show no effect on microbiota richness in comparison to DSS-treated mice (Supplementary Figure 7A). According to the beta-diversity, DINF-treated mice exhibited a significant different microbiota composition with DSS-treated or DM-treated mice (Supplementary Figure 7B). At phyla level, an increase of *Desulfobacterota* in comparison to DSS is observed (Supplementary Figure 7C). A differential analysis (DESeq2) was applied to identify differences in mucosal microbiome composition between groups. At days 14, 27 genus mainly belonging to the *Firmicutes* phylum (21/27) were significantly different (Supplementary Figure 7D). Among the *Firmicutes,* 3 species belonging to the genus of *Lachnospiraceae_NK4A136_*group (Clusters_19, 156 and 175) are underrepresented, and 1 (Cluster_136) is over-represented with the Infatrini Peptisorb® formula in comparison to Modulen IBD® (Supplementary Figure 7D). A species belonging to the *Erysipelotrichaceae* family is strongly underrepresented with Infatrini Peptisorb® formula (cluster_247, unknown genus: 20.1 log2 fold change in DINF vs DM, Supplementary Figure 7D). At the opposite, 2 genus belonging to the *Bacteroidota* phylum, family *Muribaculaceae* (Cluster_98 and 150), and 1 genus belonging to the *Actinobacteriota* phylum, family *Atopobiaceae* (Cluster_304), are increased by Infatrini Peptisorb® formula in comparison to Modulen IBD® (Supplementary Figure 7D).

### EEN with Modulen IBD® promotes epithelial cellular properties in a 3D colonic organoid model

The intestinal barrier restitution suggests an improvement of epithelial cellular functions. We investigated cell survival, proliferation, and differentiation in a 3D organoid model, performed at sacrifice days 7 or 14, and cultured for up to 9 days. The survival of organoids from DSS crypts is decreased during culture by 47 to 54%, and 39 to 46%, respectively at day 7 and day 14 (Figure 4A). The mean surface area of mature organoids is higher in DSS condition during culture at the inflammatory peak (day 7) compared to those isolated from control mice (Figure 4B). The immature organoids tend to have the same surface increase (Figures 4B and C). However, at day 14, i.e. in the recovery period, DSS mature organoids present a diminished surface area at day 3 and day 6 of culture, and an increased one at day 9 of culture (Figure 4C). The same evolution is observed for immature organoids. This could be explained by a different modulation or delay of enhanced proliferation during the remission post-inflammatory phase. Moreover, no significant difference in terme of immature versus mature structure number have been observed in control, DSS and DM conditions at days 7 and 14 (Supplementray Figure 8A). Although these DSS structures have an increased surface area during culture, the capacity to generate mature structures (budding) is decreased, with a maximum mean of budding organoids per well of 4% at day 9 of culture (Figure 4D). In addition, these DSS organoid bursts due to the excess of apoptotic cells in the lumen are considered as dead organoids (Supplementary Figure 8B). This shows that cellular properties are impaired in the DSS condition. In comparison, DM promotes cell survival, restoring it to the control levels (Figure 4A). Immature and mature organoids from crypts of DM-treated mice have similar growth to the control organoids (Figures 4B and C), and have an increased capacity to generate mature structures (budding), as seen by the budding structures at day 9 (Figure 4D). The budding structures are comparable to newly formed crypt within the organoid, suggesting a re-epithelialization capacity.

**Figure 4.**
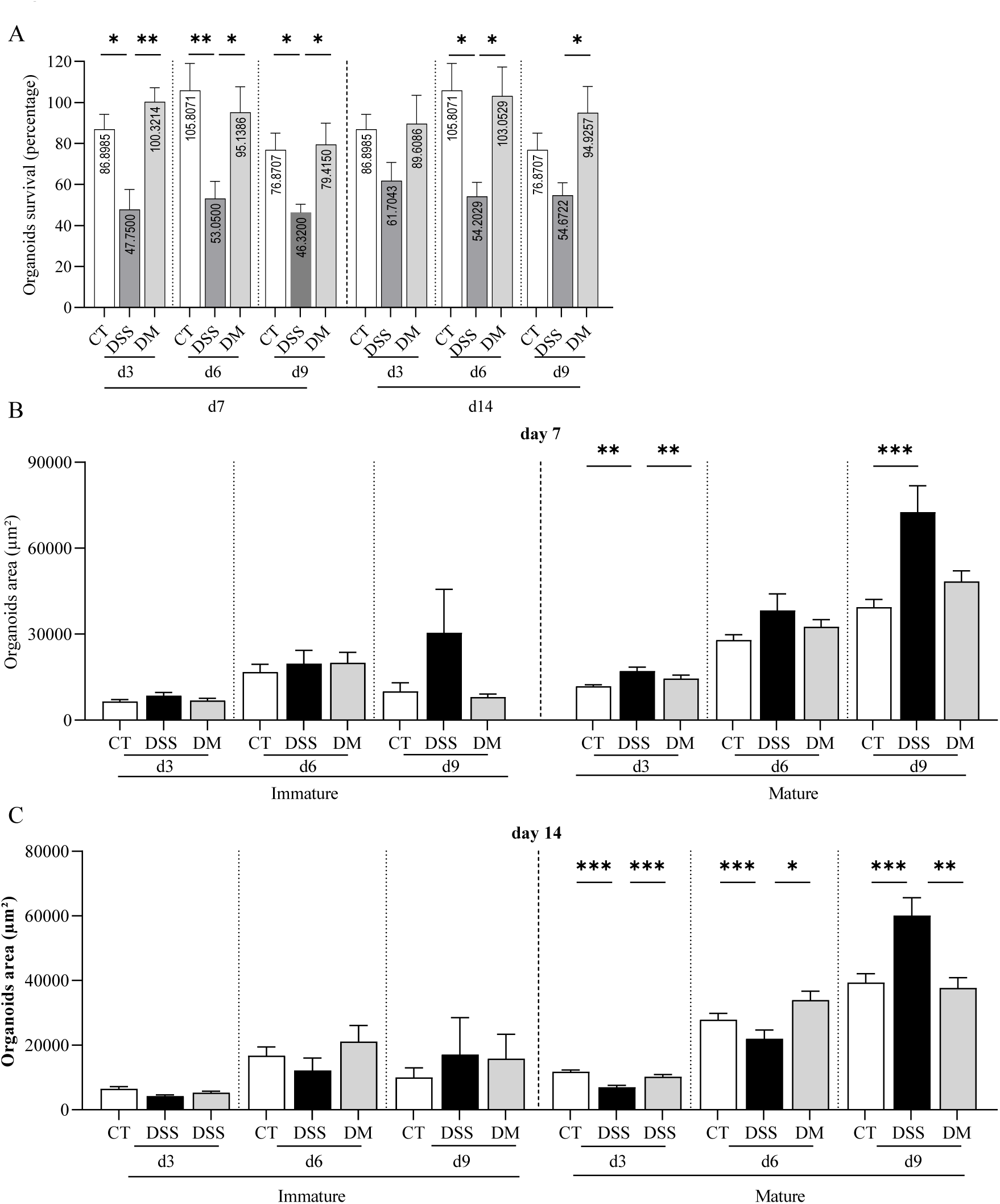
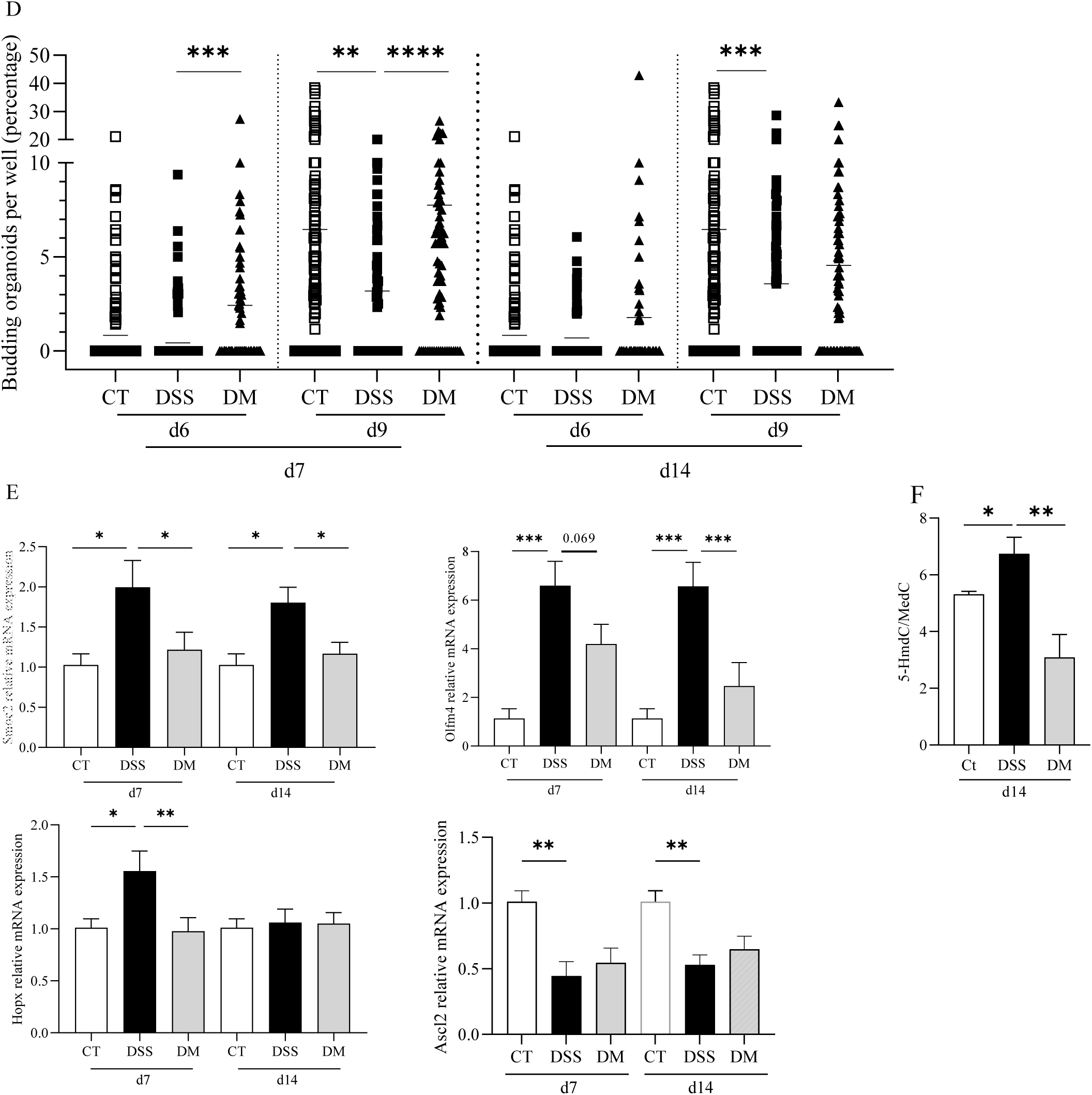
EEN promotes intestinal cellular properties in a 3D organoid model. **A:** organoid survival expressed in percentage from d3 to d9 of culture. **B:** organoids area shown in µm² of immature organoids (left), and mature organoids (right) at the inflammatory peak (d7). **C:** organoids area in µm² of immature and mature organoids at d14. **D:** budding organoids per well during culture. Only organoids with at least one bud are counted. **E:** mRNA relative expression of Smoc2, Olfm4, Hopx and Ascl2 from crypts of CT, DSS, and DM mice. **F:** percentage of methylated cytosine at position 5. F: Data are expressed as mean ± SEM; at least n>6 mice per group. *P-values*: * p<0.05; ** p<0.01; ***p<0.001; ****p<0.0001.

In addition to its impact on the ISC abilities to survive, growth and differentiate, Modulen IBD® (DM) was able to normalize the mRNA expression of SPARC-related modular calcium-binding protein 2 (*Smoc2*), a marker of crypt base columnar (CBC) stem cells at both days 7 and 14 (Figure 4E). Treatment with Modulen IBD® was also able to restore a normal mRNA expression of Olfactomedin-4 *(Olfm4)*, a marker of CBC stem cells (Figure 4E). Modulen IBD® also normalized the mRNA expression of Homeodomain-only protein X (*Hopx*), a marker of +4 stem cells at day 7 (Figure 4E). Finally, DSS treatment diminished the mRNA expression of Achaete-scute complex homolog 2 (*Ascl2*), a marker of CBC, and treatment with Modulen IBD® was unable to restore a normal expression (Figure 4E). These different cellular properties observed in a physiological *in vitro* environment suggest epigenetic modifications. As expected, the percentage of 5-methyl deoxycytosine is decreased in colonic epithelium derived from DSS treated mice and, normalized in DM conditions (Figure 4F). This suggest that Modulen IBD® may restore the methylation at position 5 of cytosine at the CpG sequences, an important epigenetic regulation. However, treatment with Modulen IBD® was unable to normalize the mRNA expression of colonocytes markers (Ap1, Sis and Fab1), Goblet cells (Clca), enteroendocrine cells (Cgc), and Tuft cells (Sucr), Mucins (Muc2, Muc4 and, Muc3), antimicrobial peptides (Tff3 and, Mmp7), tight junction protein (Ocld) and, receptor of IgA translocation (PigR) in context of colitis induced by DSS at days 7 and 14 (Supplementary Figures 9 A-M).

### DM promotes mucosal healing through gut microbiota changes

Because of the modifications of microbiota, the ability of EEN to induce its effect via the gut microbiota was studied (Figure 5). After fecal microbiota transplantation (FMT) and DSS administration, mice who have received DSS and DM FMT develop colitis, as evidenced by their weight loss and DAI (Figures 5A and B). In addition, macroscopic inflammation is revealed by the colonic shortening and thickening at day 7 and day 14 (Figures 5C and D). However, DM FMT mice tend to have decreased colonic shortening at days 7 and 14, and thickening at day 7. Macroscopic inflammation and edema scores are decreased by DM FMT at day 14, even reaching no edemas for two mice (Figures 5E and F). Also, DM FMT normalizes the intestinal permeability raised by DSS, at both sacrifice days (Figure 5G).

**Fig. 5:**
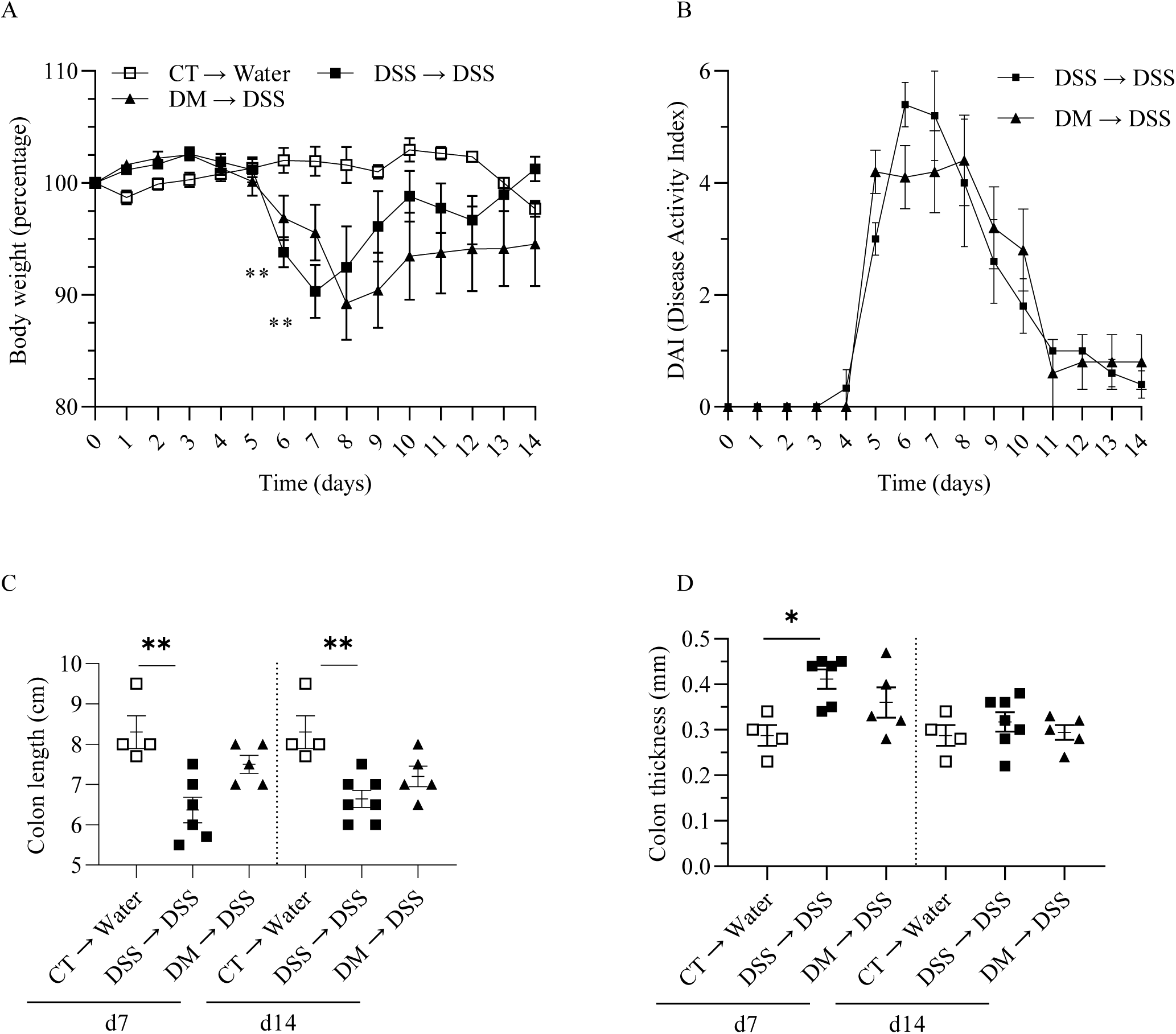
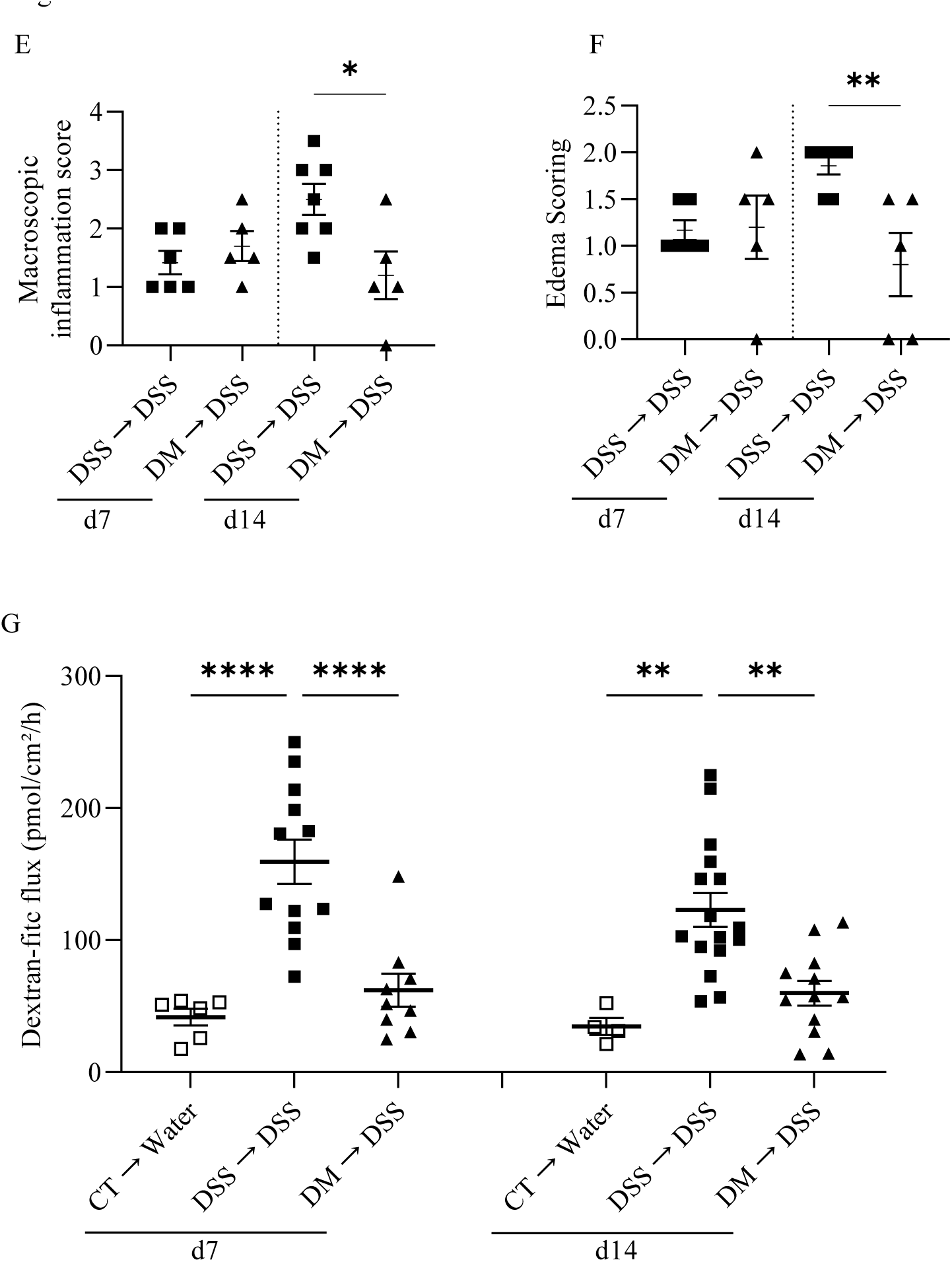
Intestinal microbiota drives EEN-induced remission. **A:** bodyweight evolution of FMT control mice (CT → water), FMT DSS mice (DSS → DSS), and FMT DM mice (DM → DSS) (legend at the top). **B:** DAI (same legend as A) (n>4). **C:** colon length in cm. **D:** colon thickness in µm. **E:** macroscopic inflammation score (n>5). **F:** edema score. G: intestinal permeability performed in duplicate, each point equivalent to a duplicate. Data are expressed as mean ± SEM; at least n>4 mice per group. *P-values*: * p<0.05; ** p<0.01; ***p<0.001; ****p<0.0001.

## DISCUSSION

EEN allows clinical remission, with mucosal healing in CD patients. Although these beneficial anti-inflammatory effects are known, the mechanisms of action of EEN remain poorly understood. Thus, the research objective was to develop an experimental model to study the mechanisms of action of EEN, along with the role of the nutritional composition, and TGF-β2 content. In our inflammatory model, DM promotes weight recovery, decreases clinical and macroscopic inflammation, attenuates microscopic inflammation, and normalizes gut permeability. Our study is the first to demonstrate these anti-inflammatory properties, as well as the epithelial restitution. In addition, we present evidence of the importance of the gut microbiota in partly mediating these beneficial effects.

Modulen IBD® significantly reduced macroscopic parameters, apart from colon length. These beneficial effects of Modulen IBD®, particularly on the weight loss and the inflammatory parameters that we observed, confirm the results obtained in a previous study carried out in a mouse model.^19^ The literature also contained studies on the anti-inflammatory properties of various EEN formula in mouse models of IBD.^7^ Nevertheless, the majority of these murine studies include the use of EEN formula that are not specific to CD, and some of them present limits in their experimental protocols. Our research was, to our knowledge, the first to put forward a robust murine inflammatory model and to address mechanisms of action of CD-specific EEN formula.

The beneficial effects of the Modulen IBD® formula were potentially related to nutritional intakes.^7^ The casein protein composition induced derivatives such as β-casofensin, which is known to reduce macroscopic and microscopic inflammation and prevent Goblet cell loss.^20^ In addition, amino acids derived from casein degradation may be involved in the remission process. For example, lysine had anti-inflammatory properties in a DSS model, and glutamic acid plays an important role in intestinal homeostasis maintenance.^21, 22^ Similarly, lipids and their derivatives could promote healing effects. Particularly, palmitic acid and medium-chain triglycerides improve cell proliferation and IgA production.^23–25^ To better understand the mechanisms of action, we used a second isocaloric formula, Infatrini Peptisorb®. Although this second formula chosen is not specifically intended for CD, its nutritional composition is very similar to Modulen IBD®. Based on nutritional intake, both formula restored the body weight of the treated mice and improved their well-being. Meanwhile, only the treatment with Modulen IBD® significantly improved macroscopic and edema scores from day 7 (inflammatory peak), and normalized intestinal permeability, suggesting a return of appropriate epithelial functionality. Nevertheless, treatment with Infatrini Peptisorb® seemed to improve colonic thickness on days 7 and 14, and the macroscopic score on day 14. This suggests an anti-inflammatory effect of the nutritional composition. By comparing the two formula compositions, the quantities of folic acid are higher in Infatrini Peptisorb®. The anti-inflammatory capabilities of folic acid were demonstrated in an *in vitro* study, which showed that folic acid changed the pro-inflammatory status of BV-2 microglial cells activated by LPS.^26^ In TNBS model, the importance of folic acid in maintaining Foxp3+ Tregs in the colon was demonstrated, as well as the anti-apoptotic molecules Bcl2 and Bcl-xl, thus preventing colitis.^27^ In humans, a comparative study demonstrated that folic acid supplementation reduced CRP in patients with various pathologies (diabetes, obesity, metabolic syndrome, etc.).^28^ Another difference between the two EEN formula chosen is that Modulen IBD® is a polymeric formula, i.e. containing proteins in a complete form while Infatrini Peptisorb® is a semi-elemental formula, in which these proteins are partly hydrolyzed. A study by Souza et al. suggests that an elemental formula is less anti-inflammatory than a polymeric formula, and even exacerbates colitis in a DSS model.^29^ However, the nutritional composition of their formula differs from those we used: 16.7% lipids, 64% carbohydrates, and 19.3% nitrogen source (proteins). Moreover, the Infatrini Peptisorb® formula is semi-elemental, thus the proteins are not fully hydrolyzed. Finally, clinical data indicate that clinical remission of patients is not affected by the type of formula used.^30^ Therefore, these components do not explain the better results obtained with Modulen IBD®.

We subsequently investigated the involvement of TGF-β2 contained in Modulen IBD®, in contrast to the Infatrini Peptisorb® formula in which it was not detected. The relevance of this cytokine in IBD has been previously demonstrated.^31, 32^ In a DSS model, deletion of the TGF-β receptor induces exacerbation of colitis.^32^ Furthermore, Miyoshi et al. demonstrated that TGF-β in the extracellular matrix promotes colonic crypt regeneration dependent on Wnt5a, a ligand produced by mesenchymal cells.^31^ Concerning TGF-β levels in patient tissues, one study showed no difference in TGF-β1 expression between colonic mucosa of IBD patients and healthy subjects.^33^ At the level of the *lamina propria*, TGF-β2 and TGF-β3 expressions are increased in relapsed IBD patients^32^. Similarly, the intestinal mucosa of IBD patients has a defect in the TGF-β signaling pathway with deregulations of Smads,^11, 34^ i.e. a decreased phosphorylation of Smad3 in the *lamina propria*, and increased Smad7 in the mucosa and *lamina propria* of IBD patients.^11, 35^ Since Smad7 is an inhibitor of the TGF-β signaling pathway, this blocks its anti-inflammatory effects. This signaling pathway is reestablished after the inhibition of Smad7.^35^ Thus, the TGF-β2 richness of Modulen IBD® could explain the barrier restitution obtained in our model, unlike Infatrini Peptisorb®. Our results confirm those of a previous study carried out on another mouse model of colitis (*IL10*^−/−^).^36^ Nevertheless, our data are the first to present more detailed results on the components of the intestinal barrier and ISCs capabilities. Moreover, as demonstrated by supplementation of Infatrini Peptisorb® with TGF-β2, our work highlights the involvement of TGF-β2 on colonic thickness, macroscopic, and edema scores, as well as normalization of colonic permeability. These effects on macroscopic and microscopic parameters demonstrate that TGF-β2 potentiates the nutritional composition. Moreover, considering the loss of these effects after neutralization of TGF-β2 in Modulen IBD®, we demonstrate the importance of this cytokine in mucosal healing and functionality during EEN with Modulen IBD®. This phenomenon was notably observed in a recent clinical study, where only patients treated with Modulen IBD® had an improvement on histological inflammation.^37^ Also, supplementation of Infatrini Peptisorb® with TGF-β2 shows a similar level of resolution as Modulen IBD®, notably with the decrease of edema and normalization of intestinal permeability. Moreover, the effects obtained with TGF-β are more accentuated at day 14, i.e. at the recovery phase, suggesting the importance of this cytokine for cellular regeneration.

In addition to direct effects of nutrients and TGF-β, a change in the microbiota induced by Modulen IBD® could participate in the beneficial effects of this formula on mucosal healing. Thus, we sequenced the gut microbiota to study variations in microbial populations. Of note, microbiota analyzes correspond to the mucosal microbiota and not the fecal microbiota, allowing understanding which bacterial populations could have an effect on the epithelium. At the inflammatory peak (d7), *Bacteroidota*, *Proteobacteria* and *Desulfobacterota* are increased in DM in comparison to DSS, while *Firmicutes* are decreased. Nevertheless, within the *Firmicutes* phylum, some species are decreased, notably *Clostridia* at d7, while others are increased. For example, in our results, despite a decrease in *Firmicutes* after EEN, we found an increase of *Lachnospiraceae* within this phylum, which is also reported by clinical studies.^38^ This family produces indole-type metabolites involved in tissue repair and maintenance of homeostasis.^39^ Moreover, alpha-diversity seems to be improved after treatment in our model.

Subsequently, we investigated the involvement of modified gut microbiota in the anti-inflammatory and pro-regenerative properties of TGF-β2-rich Modulen IBD®. We demonstrated that these beneficial properties are mediated in part by the intestinal microbiota, notably by the decrease of macroscopic and edema scores, and the normalization of colonic permeability. The role of the microbiota suggests that its maintenance in patients might be a possibility to maintain remission. It has been shown that the gut microbiota of patients returned to its pre-treatment composition once EEN was completed, because of the reintroduction of solid food.^40^ This supports some of the proposals made to sustain this post-treatment microbiota, such as through the administration of prebiotics and probiotics. However, this would require a better understanding of the gut microbiota and its dynamics upon EEN in IBD patients. In addition, studies would be helpful to understand whether this addition to EEN would not impair remission.

Finally, we studied the cellular capacities of the colonic crypts, treated or not with Modulen IBD®. The 3D organoid model was used to study the gut epithelium and its capacities independently of the other components of the intestinal barrier. We found that organoid survival was decreased under inflammatory conditions, reflecting the inability of DSS-treated crypts to close and form mini-intestines. Similarly, these organoids showed increased proliferation, which may be caused by the alteration of cellular pathways of proliferation and differentiation, demonstrated also by the decrease in neo-crypt formation. These parameters were improved in organoids developed from DM mice. In particular, the formation of neo-crypts was increased, suggesting an enhanced capacity of re-epithelialization of these cells, thus an increased ability to induce mucosal functional healing. The culture conditions of organoids, with the presence of the epithelium only, indicated that Modulen IBD® probably induced a genomic cellular imprinting. Our results showed an improvement of cytosine methylation at position 5, a DNA methylation marker, after Modulen IBD® in a comparable way to controls. DNA methylation is altered in IBD patients, and appears to be involved in their pathophysiology,^41, 42^ and in mice, DSS altered epigenetic in a microbiota-dependent way.^43^ Epigenetics deserves to be studied in murine and clinical studies to better understand the mechanisms of EEN, as shown by our results that highlight for the first time the possible epigenetic modifications induced by EEN, including Modulen IBD®, in IBD.

In conclusion, our data demonstrate the anti-inflammatory and pro-regenerative properties of EEN treatment with Modulen IBD®. Although nutritional composition appears to be involved in these properties, TGF-β2 plays a major role in restoring the barrier function. In addition, the resulting gut microbiota is involved in the maintenance of EEN-induced homeostasis. Finally, cellular pro-regenerative capacities are improved, probably through genomic imprinting. Our results are the first to provide details on the mechanisms by which EEN promotes the epithelium restitution in a mouse model of CD. Although this is a significant discovery, many questions remain and require future murine and human studies. This continuation will be necessary to improve therapeutic strategies and management of CD patients.

## Acknowledgments

Thanks go to Céline Barde, dietitian at the children hospital of Purpan, for discussion and counselling on Modulen IBD® and Infatrini Peptirosb®. We are grateful to the INRAE MIGALE bioinformatics facility (MIGALE, INRAE, 2020. Migale bioinformatics Facility, doi: 10.15454/1.5572390655343293E12) for providing computing and storage resources. Thanks to the Metagenomic16S service for microbiota analyses (CRI, Paris, France). Also, thanks to Sophie Allart and Simon Lachambre for their technical assistance at the cellular imaging facility of INSERM UMR 1291, Toulouse. Thanks to Timothé Durand-Plavis and Samantha Milia for their technical assistance at the experimental histopathology facility of the INSERM / UPS US006 CREFRE, Toulouse Purpan, France. Thanks to Frédéric Martins for Fluidigm® lecture. Finally, thanks to Dr. Marie Carrière for methylation analysis.

