## supplemental figures and tables for "Promoting intestinal healing by exclusive enteral nutrition with TGF-β in a mouse model of colitis"

Supplementary Figure 1

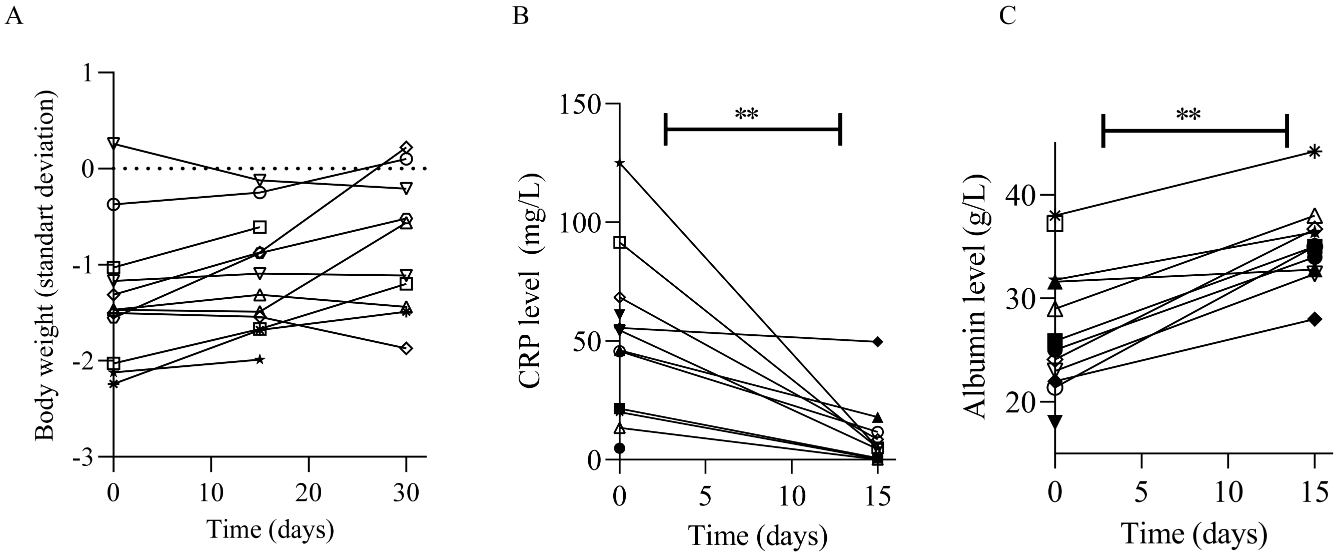

Supplementary Figure 2

A

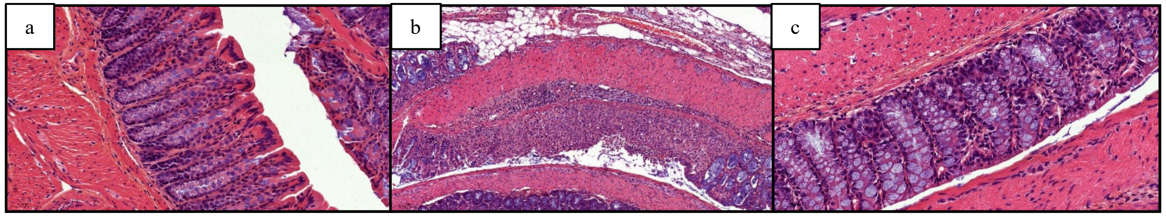


B

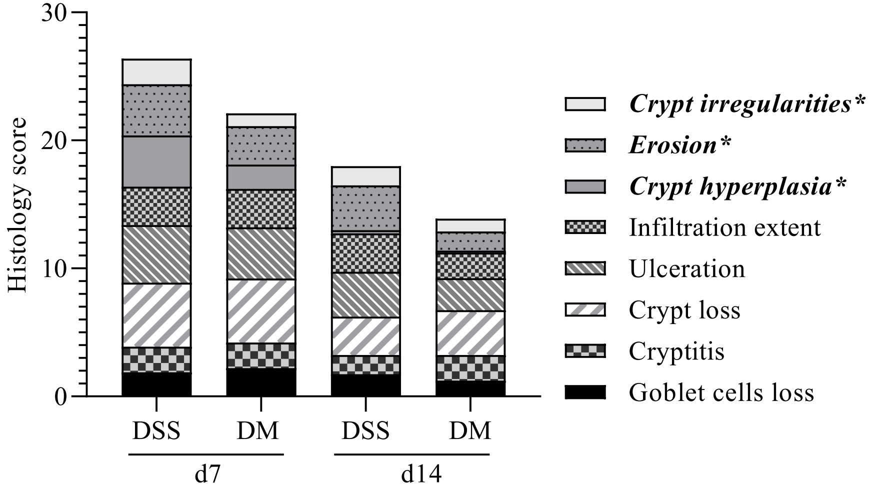

Supplementary Figure 3

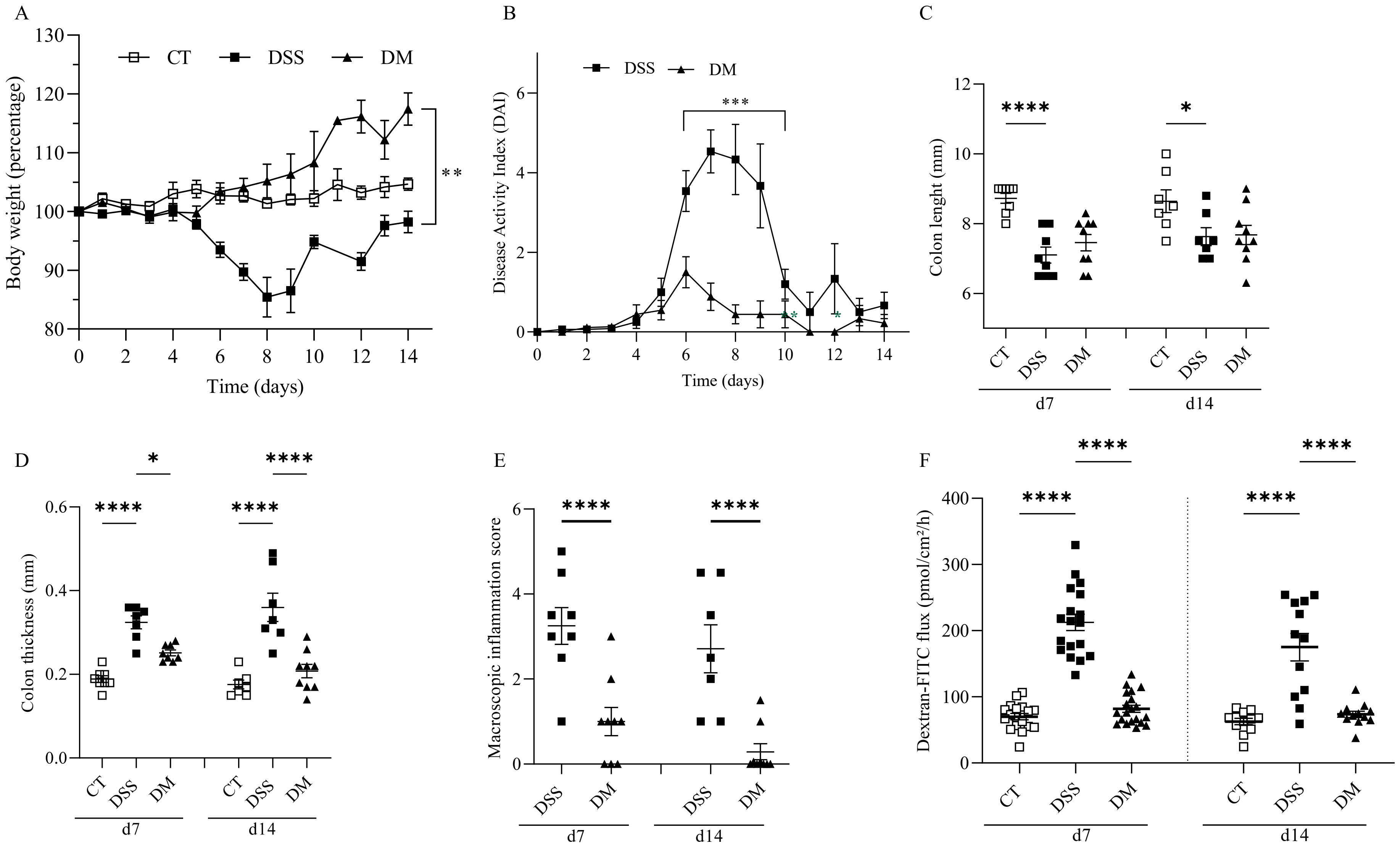

Supplementary Figure 4

A

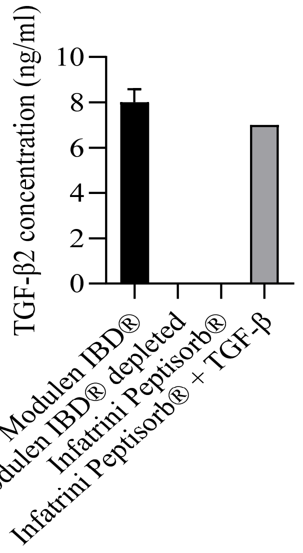

Supplementary Figure 5

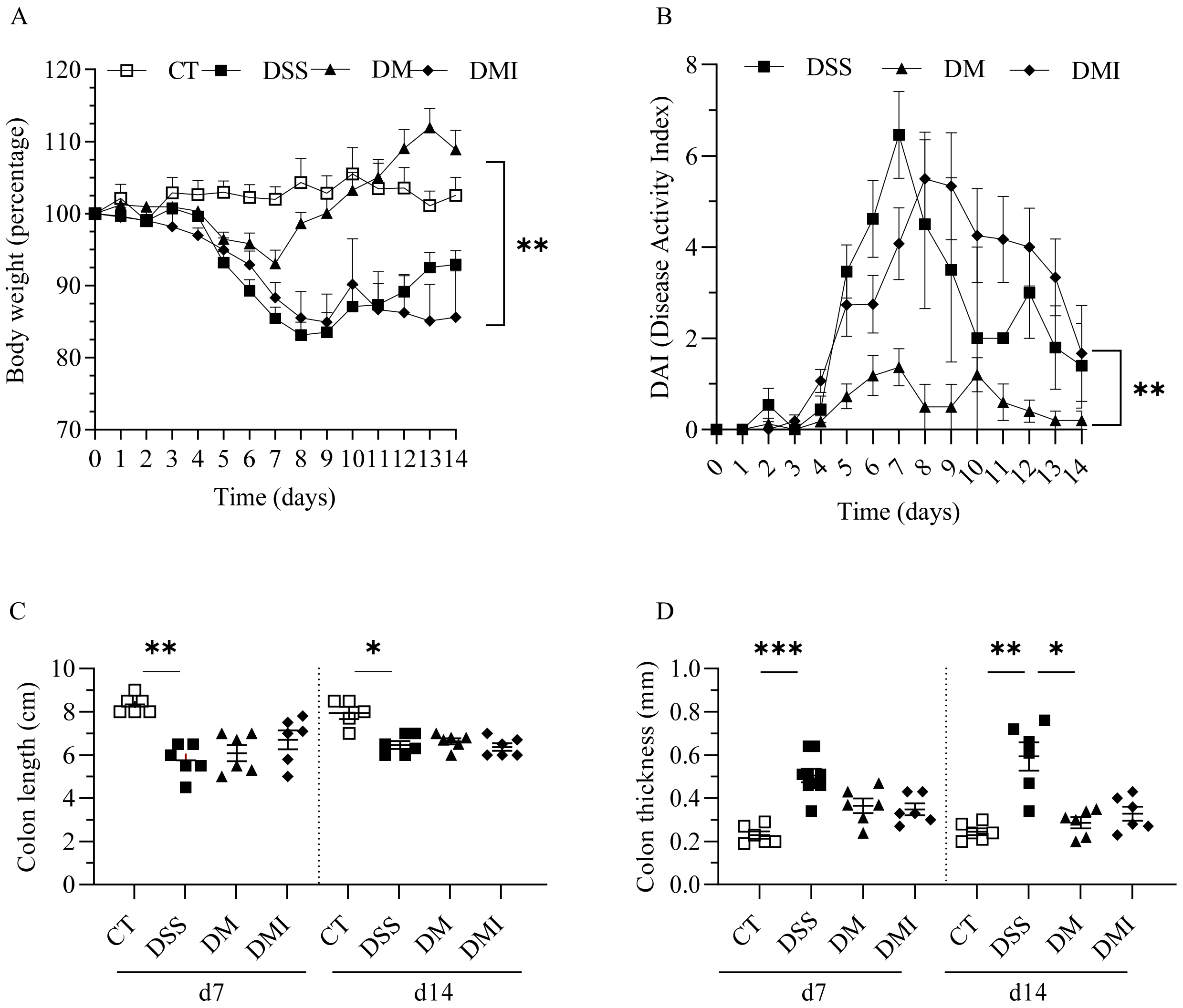

Supplementary Figure 5

E

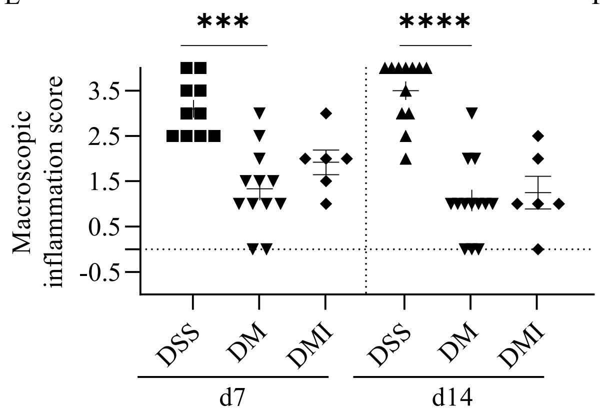


F

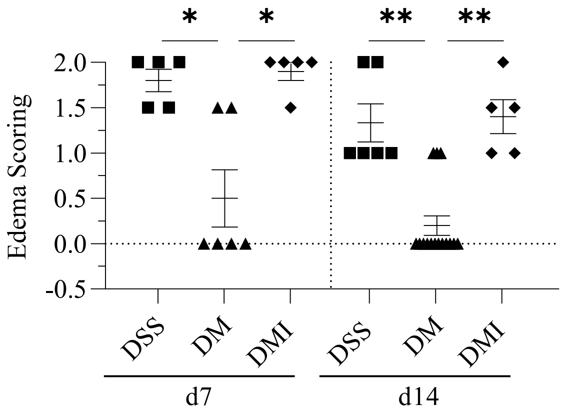


G

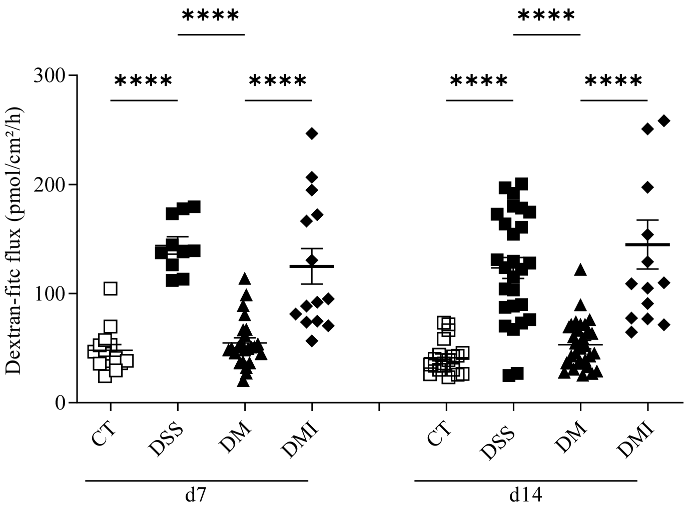

Supplementary Figure 6

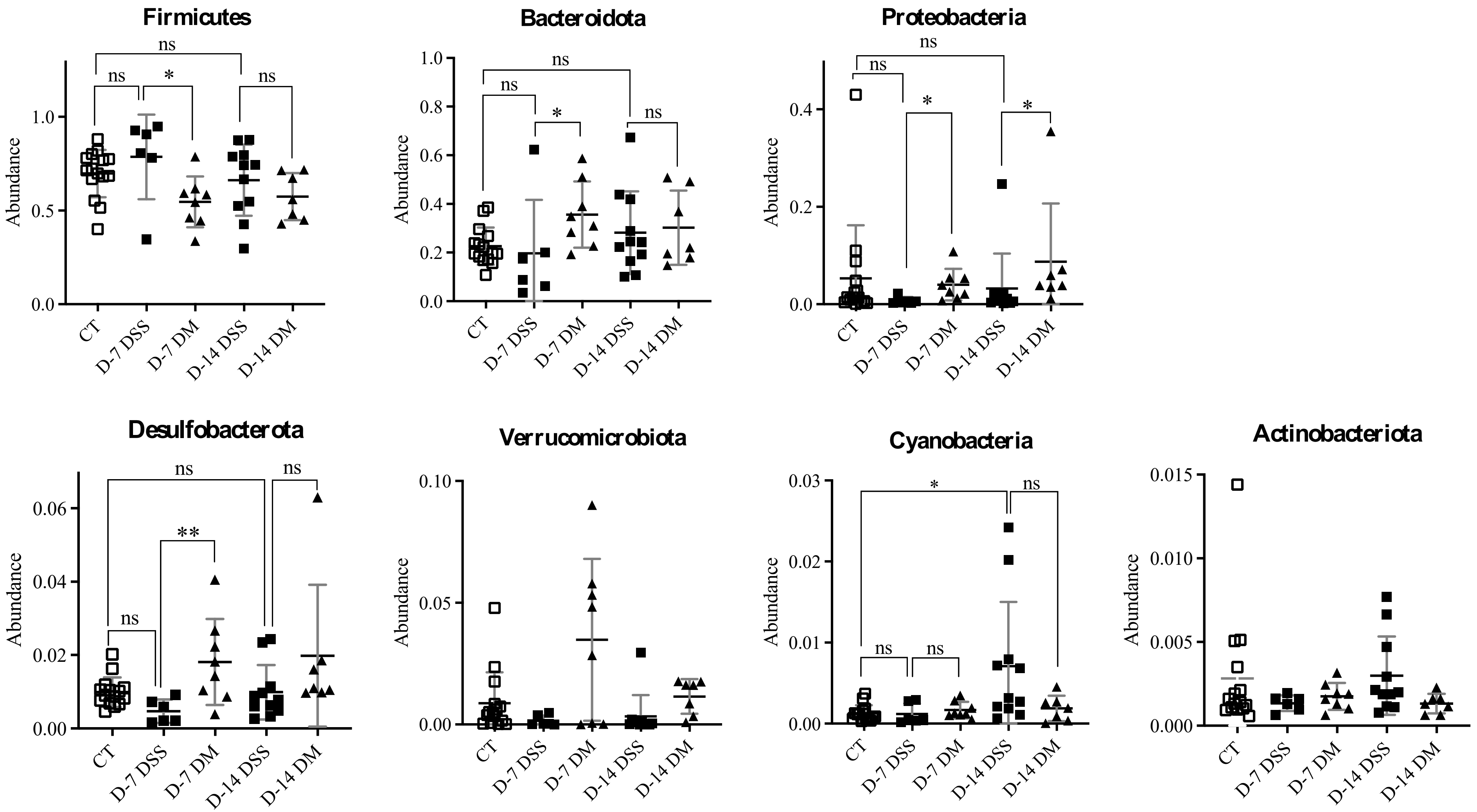

Supplementary Figure 7

A

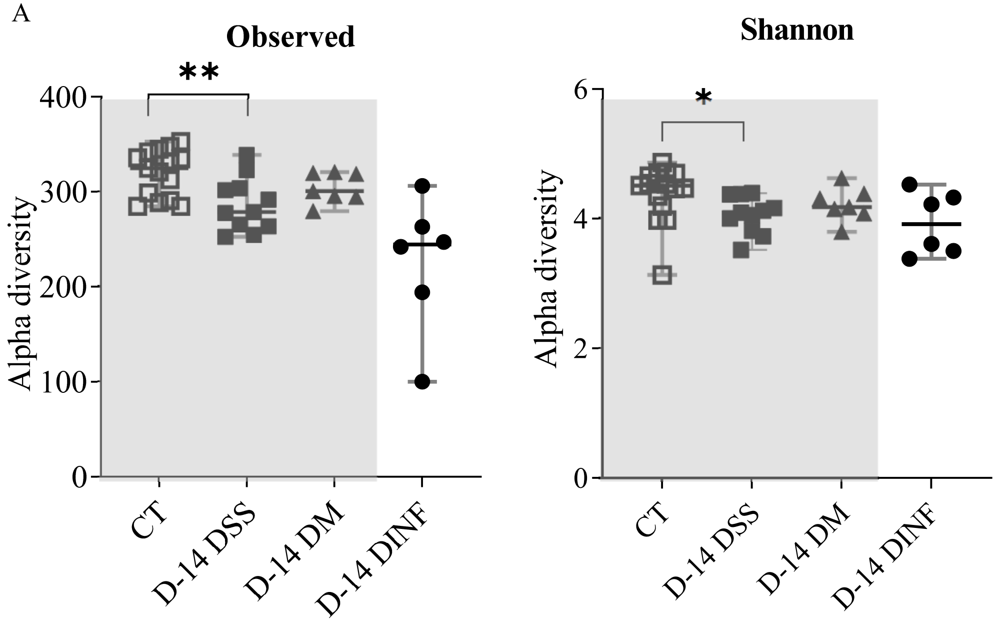


B

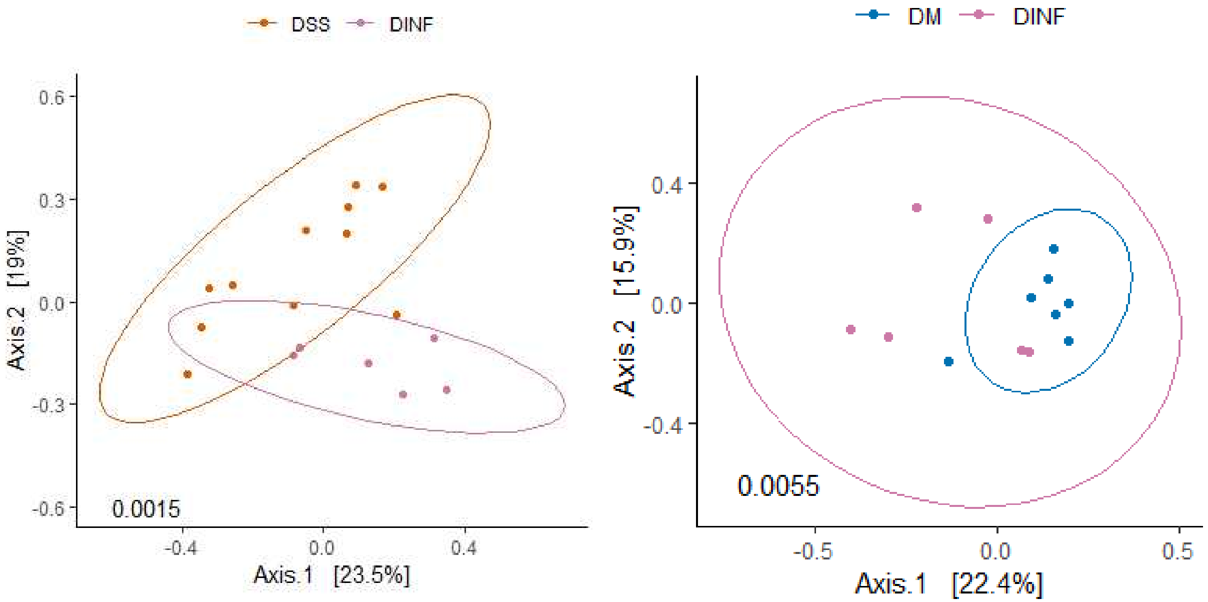


C

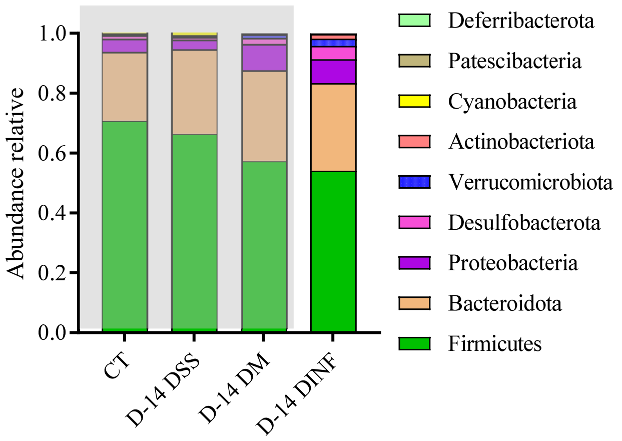

Supplementary Figure 7

D

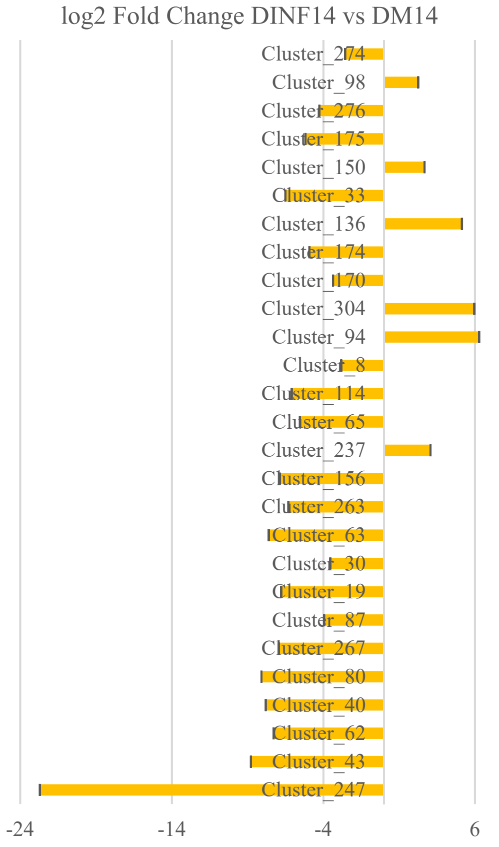


| OTU | Phylum / Class / Order / Family / Genus |
|---|---|
| Cluster_247 | Firmicutes / Bacilli / Erysipelotrichales / Erysipelotrichaceae / unknown genus |
| Cluster_43 | Proteobacteria / Gammaproteobacteria / Enterobacterales / Enterobacteriaceae / Lelliottia |
| Cluster_62 | Firmicutes / Clostridia / Lachnospirales / Lachnospiraceae / unknown genus |
| Cluster_40 | Firmicutes / Bacilli / Acholeplasmatales / Acholeplasmataceae / Anaeroplasma |
| Cluster_80 | Firmicutes / Clostridia / Lachnospirales / Lachnospiraceae / Lachnoclostridium |
| Cluster_267 | Firmicutes / Clostridia / Clostridiales / Clostridiaceae / Clostridium sensu stricto 1 |
| Cluster_87 | Bacteroidota / Bacteroidia / Bacteroidales / Marinifilaceae / Odoribacter |
| Cluster_19 | Firmicutes / Clostridia / Lachnospirales / Lachnospiraceae / Lachnospiraceae NK4A136 group |
| Cluster_30 | Firmicutes / Clostridia / Lachnospirales / Lachnospiraceae / Lachnoclostridium |
| Cluster_63 | Firmicutes / Clostridia / Lachnospirales / Lachnospiraceae / unknown genus |
| Cluster_263 | Firmicutes / Clostridia / Lachnospirales / Lachnospiraceae / [Eubacterium] xylanophilum group |
| Cluster_156 | Firmicutes / Clostridia / Lachnospirales / Lachnospiraceae / Lachnospiraceae NK4A136 group |
| Cluster_237 | Firmicutes / Clostridia / Lachnospirales / Lachnospiraceae / unknown genus |
| Cluster_65 | Firmicutes / Clostridia / Lachnospirales / Lachnospiraceae / unknown genus |
| Cluster_114 | Firmicutes / Clostridia / Lachnospirales / Lachnospiraceae / Roseburia |
| Cluster_8 | Firmicutes / Bacilli / Lactobacillales / Lactobacillaceae / Lactobacillus |
| Cluster_94 | Actinobacteriota / Coriobacteriia / Coriobacteriales / Atopobiaceae / Olsenella |
| Cluster_304 | Firmicutes / Clostridia / Lachnospirales / Lachnospiraceae / unknown genus |
| Cluster_170 | Firmicutes / Clostridia / Oscillospirales / Oscillospiraceae / Oscillibacter |
| Cluster_174 | Firmicutes / Clostridia / Lachnospirales / Lachnospiraceae / unknown genus |
| Cluster_136 | Firmicutes / Clostridia / Lachnospirales / Lachnospiraceae / Lachnospiraceae NK4A136 group |
| Cluster_33 | Firmicutes / Clostridia / Oscillospirales / Ruminococcaceae / [Eubacterium] siraeum group |
| Cluster_150 | Bacteroidota / Bacteroidia / Bacteroidales / Muribaculaceae / unknown genus |
| Cluster_175 | Firmicutes / Clostridia / Lachnospirales / Lachnospiraceae / Lachnospiraceae NK4A136 group |
| Cluster_276 | Proteobacteria / Gammaproteobacteria / Pseudomonadales / Pseudomonadaceae / Pseudomonas |
| Cluster_98 | Bacteroidota / Bacteroidia / Bacteroidales / Muribaculaceae / unknown genus |
| Cluster_274 | Firmicutes / Clostridia / Oscillospirales / Ruminococcaceae / unknown genus |

Supplementary Figure 8

A

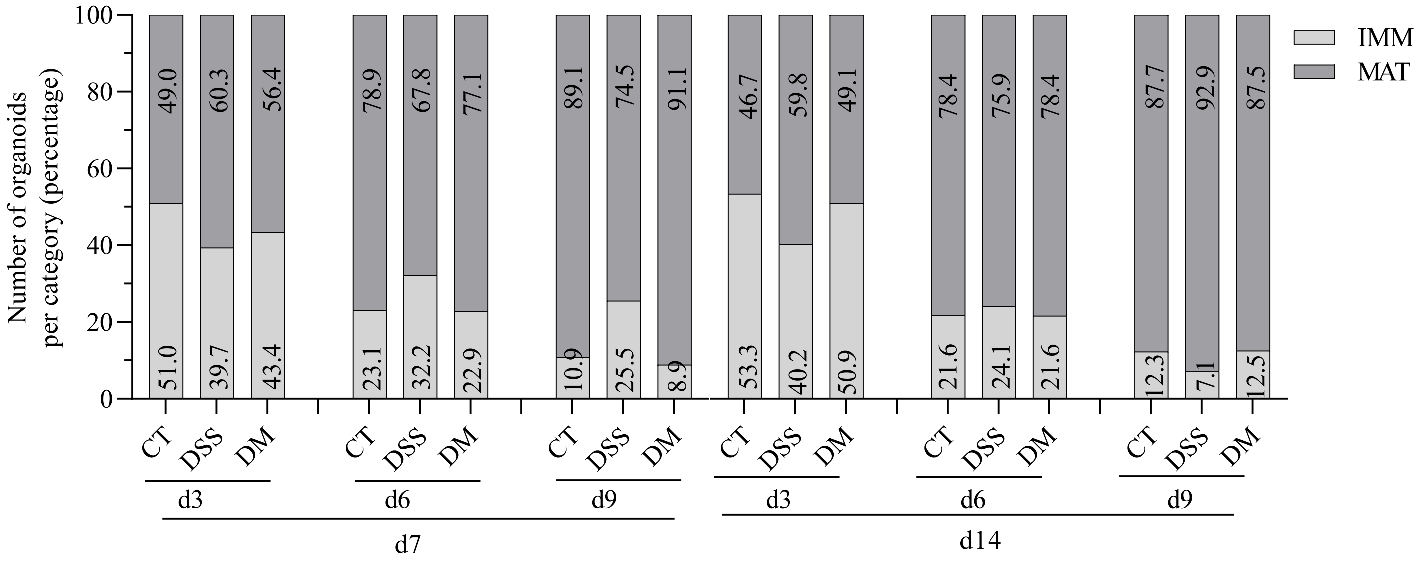


B

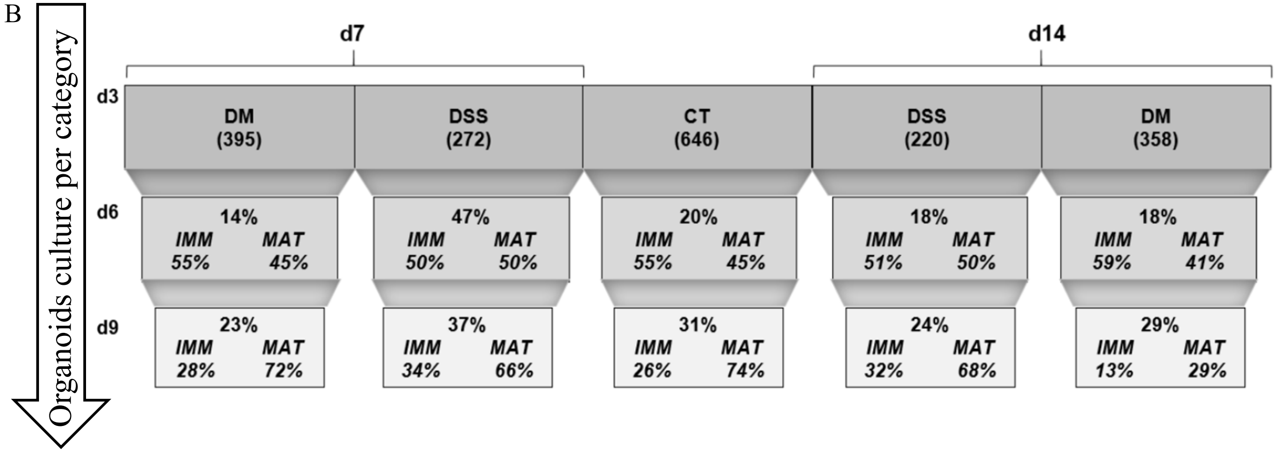

Supplementary Figure 9

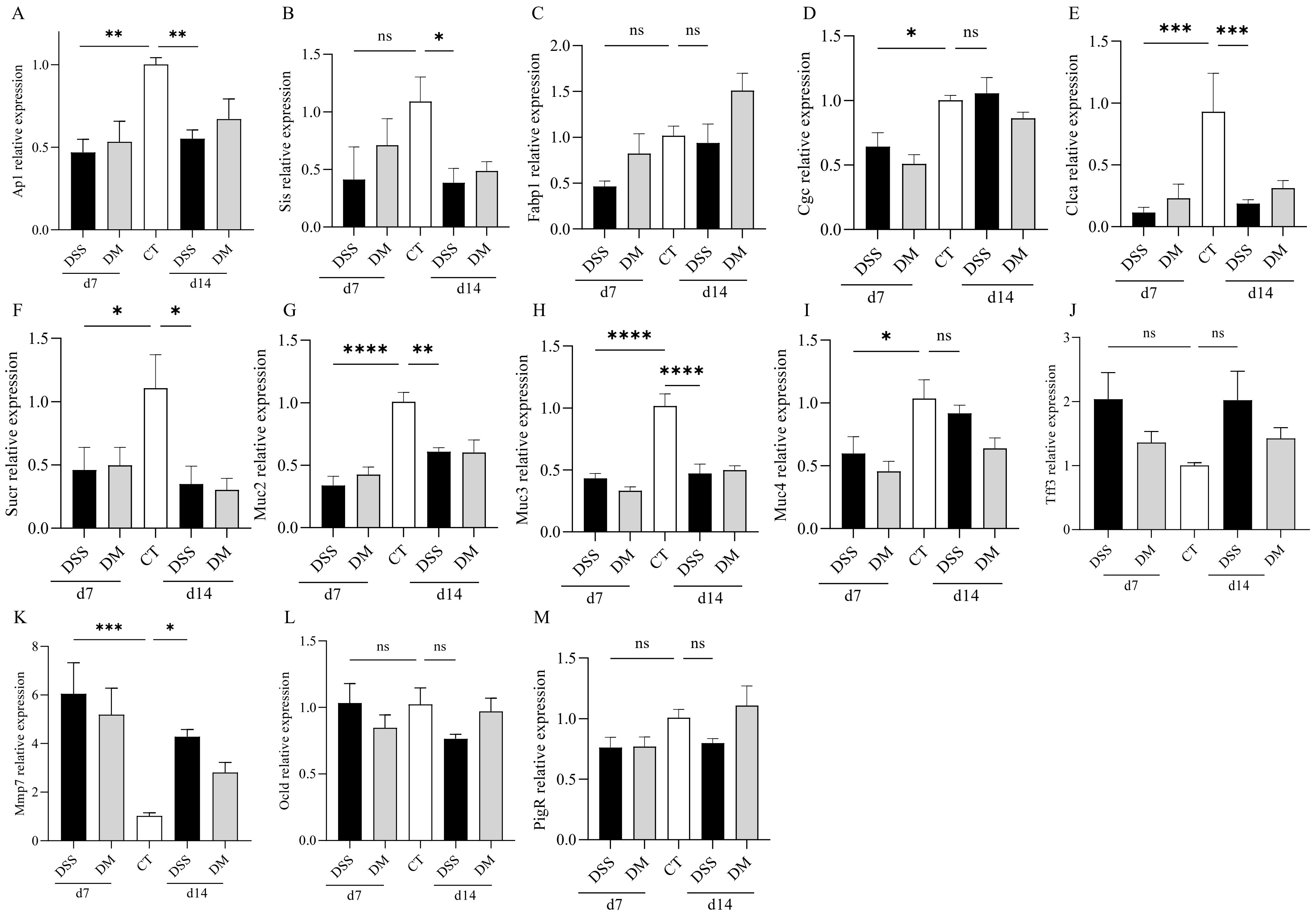

Supplementary Table 1

| Name | Forward sequence | Reverse sequence |
|---|---|---|
| Gapdh | AGGTCGGTGTGAACGGATTTG | TGTAGACCATGTAGTTGAGGTCA |
| Cxcl8 | AGCCACACTCAAGAATGGTC | GTCAGAAGCCAGCGTTCAC |
| Ascl2 | GCCTACTCGTCGGAGGAA | CCAACTGGAAAAGTCAAGCA |
| Sis | GTGCCTGCTGTGGAAGAAGTA | AACAGCCACGCTCTTCACAT |
| Fabp1 | GGGGAAAAAGTCAAGGCAGTC | TGTCGCCCAATGTCATGGT |
| Clca | GCCGGAAGATGAAGCCCTC | TTGATACAAGGACATCAGCGTTTTT |
| Muc2 | TGACTGCCGAGACTCCTACA | CTGTAGTGTGGGGTGCTGAC |
| Muc4 | TGGCTACAAAGGCTACCACC | CCCTCACTACATGGGGACAC |
| Muc3 | CTCTGCCTGCTGGCTTTCAT | CTGTTTTCCCCGTCTGTGGTT |
| Cgc | TACACCTGTTCGCAGCTCAG | GCTGGGAATGATCTGGGGTT |
| SucnR | GCTCCTGGCAGAGTTTTCTGT | TGCAGAAGAGGTAGCCAAACA |

Supplementary Table 2

| | Per 100 ml (1.0 Kcal/ml) | | |
|---|---|---|---|
| | Modulen IBD® | Infatrini Peptisorb® | % |
| **Energy (Kcal)** | 98.6 | 100 | 101 |
| **Carbohydrates (g)** | 11 | 10.2 | 93 |
| **Proteins (g)** | 3.5 | 2.6 | 74 |
| **Fats (g)** | 4.6 | 5.4 | 117 |
| **SFAs (g)** | 2.6 | 3.8 | 146 |
| **MCTs (g)** | 1.2 | 2.8 | 233 |
| **MFAs (g)** | 0.78 | 0.77 | 99 |
| **PFAs (g)** | 0.50 | 0.86 | 172 |
| **α linolenic acid (mg)** | 40 | 70 | 175 |
| **Linoleic acid (mg)** | 420 | 690 | 164 |
| **Vitamins** | | | |
| **A (µg)** | 82 | 88 | 107 |
| **D (µg)** | 0.98 | 2.4 | 245 |
| **E (mg)** | 1.3 | 2.1 | 162 |
| **K (µg)** | 5.4 | 6.7 | 124 |
| **C (mg)** | 9.4 | 14 | 149 |
| **Thiamin (mg)** | 0.12 | 0.15 | 125 |
| **Riboflavin (mg)** | 0.13 | 0.2 | 154 |
| **Niacin (mg)** | 1.2 | 0.8 | 67 |
| **B6 (mg)** | 0.17 | 0.11 | 65 |
| **Folic acid (µg)** | 24 | 16 | 67 |
| **B12 (µg)** | 0.32 | 0.3 | 94 |
| **Biotin (µg)** | 3.2 | 4 | 125 |
| **PA (mg)** | 0.48 | 0.8 | 167 |

| | Per 100 ml (1.0 Kcal/ml) | | |
|---|---|---|---|
| | Modulen IBD® | Infatrini Peptisorb® | % |
| **Sodium (mg)** | 34 | 37 | 109 |
| **Potassium (mg)** | 120 | 111 | 93 |
| **Chloride (mg)** | 73 | 75 | 103 |
| **Calcium (mg)** | 89 | 90 | 101 |
| **Phosphorus (mg)** | 60 | 45 | 75 |
| **Magnesium (mg)** | 20 | 9 | 45 |
| **Iron (mg)** | 1.1 | 1,2 | 109 |
| **Zinc (mg)** | 0.94 | 0.8 | 85 |
| **Copper (mg)** | 0.098 | 0.075 | 77 |
| **Manganese (mg)** | 0.20 | 0.006 | 3 |
| **Fluoride (mg)** | | - | |
| **Selenium (µg)** | 3.4 | 3.75 | 110 |
| **Chromium (µg)** | 5 | 4 | 80 |
| **Molybdenum (µg)** | 7.4 | 6 | 81 |
| **Iodine (µg)** | 9.8 | 19 | 194 |
| **Other nutrients** | | | |
| **Choline (mg)** | 7 | 32.1 | 459 |
| **Carnitine (mg)** | - | 2 | |
| **Taurine (mg)** | - | 7 | |
| **Inositol (mg)** | - | 25 | |
| **Osmolarity (mOsm/L)** | 290 | 295 | 102 |

SFAs : Saturated Fatty acids; MCTs: Medium; chain triglycerides; MFAs: Monounsaturated fatty acids; PFAs : Polyunsaturated fatty acids; PA: Pantothenic acid
